# High affinity enhancer-promoter interactions can bypass CTCF/cohesin-mediated insulation and contribute to phenotypic robustness

**DOI:** 10.1101/2021.12.30.474562

**Authors:** Shreeta Chakraborty, Nina Kopitchinski, Ariel Eraso, Parirokh Awasthi, Raj Chari, Pedro P Rocha

## Abstract

Transcriptional control by distal enhancers is an integral feature of gene regulation. To understand how enhancer-promoter interactions arise and assess the impact of disrupting 3D chromatin structure on gene expression, we generated an allelic series of mouse mutants that perturb the physical structure of the *Sox2* locus. We show that in the epiblast and in neuronal tissues, CTCF-mediated loops are neither required for the interaction of the Sox2 promoter with distal enhancers, nor for its expression. Insertion of various combinations of CTCF motifs between *Sox2* and its distal enhancers generated ectopic loops with varying degrees of insulation that directly correlated with reduced transcriptional output. Yet, even the mutants exhibiting the strongest insulation, with six CTCF motifs in divergent orientation, could not fully abolish activation by distal enhancers, and failed to disrupt implantation and neurogenesis. In contrast, cells of the anterior foregut were more susceptible to chromatin structure disruption with no detectable SOX2 expression in mutants with the strongest CTCF-mediated boundaries. These animals phenocopied loss of SOX2 in the anterior foregut, failed to separate trachea from esophagus and died perinatally. We propose that baseline transcription levels and enhancer density may influence the tissue-specific ability of distal enhancers to overcome physical barriers and maintain faithful gene expression. Our work suggests that high affinity enhancer-promoter interactions that can overcome chromosomal structural perturbations, play an essential role in maintaining phenotypic robustness.

## INTRODUCTION

Enhancers can control precise spatial and temporal gene expression across large genomic distances. The nuclear organization mechanisms that guide enhancers to their target promoters while preventing off-target gene activation are instrumental for establishment of transcriptional networks guiding cell fate decisions (de Laat and Duboule, 2013; Long et al., 2016). For example, genome folding by the combined action of the cohesin complex and CTCF has been suggested to facilitate enhancer recruitment when CTCF binding occurs proximal to promoters (Kubo et al., 2021; Schuijers et al., 2018; Tang et al., 2015). CTCF is thought to block extrusion of interphase chromatin by cohesin in an orientation-dependent manner, leading to close physical association of loci with convergent CTCF motifs, which become the anchors of a DNA loop (Fudenberg et al., 2016; Li et al., 2020; Nora et al., 2017; Rao et al., 2017; Schwarzer et al., 2017). CTCF-independent mechanisms that convey regulatory information from distal enhancers to promoters are less well defined but include molecular bridges such as LDB1, YY1, and the mediator complex. In addition, enhancer-promoter interactions can be induced by the act of transcription itself and spatial clustering of chromatin regions bound by the same transcription factors, chromatin regulators and with similar histone modifications (Furlong and Levine, 2018; Oudelaar and Higgs, 2021; Pombo and Dillon, 2015; Schoenfelder and Fraser, 2019).

CTCF/cohesin-mediated loops have also been shown to restrict enhancer-promoter interactions across its CTCF-bound anchors. Indeed, CTCF motifs are frequently found at the boundaries of topologically associated domains (TADs)— defined as regions of high self-interaction insulated from their genomic neighbors—and most enhancer-promoter pairs are found within the same domain (Bonev et al., 2017; Dixon et al., 2012; Nora et al., 2012; Rao et al., 2014; Sexton et al., 2012). Many of the critical insights gained in the quest to define the impact of nuclear organization on gene expression have come from the genetic manipulation of domains containing genes important for mammalian development. For example, these studies have shown that boundary deletion can expose promoters to neighboring enhancers, causing ectopic gene activation and developmental phenotypes (Kraft et al., 2019; Lupianez et al., 2015; Narendra et al., 2015). At other loci however, domain disruption has only modest or no measurable impact on gene expression and animal physiology (Amandio et al., 2021; Despang et al., 2019; Ghavi-Helm et al., 2019; Rodriguez-Carballo et al., 2020; Williamson et al., 2019). These opposing observations suggest the existence of locus and tissue-specific mechanisms by which enhancers regulate promoters that may not rely on CTCF-mediated loops. It also highlights our incomplete understanding of how the interplay of enhancer-promoter interactions and CTCF-mediated loops contribute to faithful gene expression (Beagan and Phillips-Cremins, 2020; Misteli, 2020).

The transcription factor SOX2 is essential for establishment of pluripotent states in cultured embryonic stem (ES) cells as well as in the epiblast of pre-implantation embryos (Avilion et al., 2003). SOX2 is also critical for the differentiation of other epithelial lineages such as those in the nervous system, retina, and anterior foregut derivatives (trachea, lungs, esophagus, and stomach) (Favaro et al., 2009; Que et al., 2007; Taranova et al., 2006). The murine *Sox2* locus is found in a gene desert, which has facilitated the identification of the enhancers responsible for its tissue-specific expression (Uchikawa and Kondoh, 2016). A cluster of CTCF motifs is found adjacent to the *Sox2* promoter and therefore interfering with CTCF binding enables the study of how CTCF/cohesin-mediated loops interplay with other mechanisms to influence enhancer recruitment (Fig. 1A). These characteristics have made *Sox2* an ideal testbed for functional studies of gene regulation that have, for example, characterized the direct influence of enhancer distance and CTCF-mediated loops on transcription (Alexander et al., 2019; Huang et al., 2021; Zuin et al., 2021). Surprisingly, despite its frequent use in studies of gene regulation in vitro, we still do not know how the different mechanisms of enhancer recruitment affect *Sox2* expression in vivo and ultimately its developmental functions.

**Figure 1:**
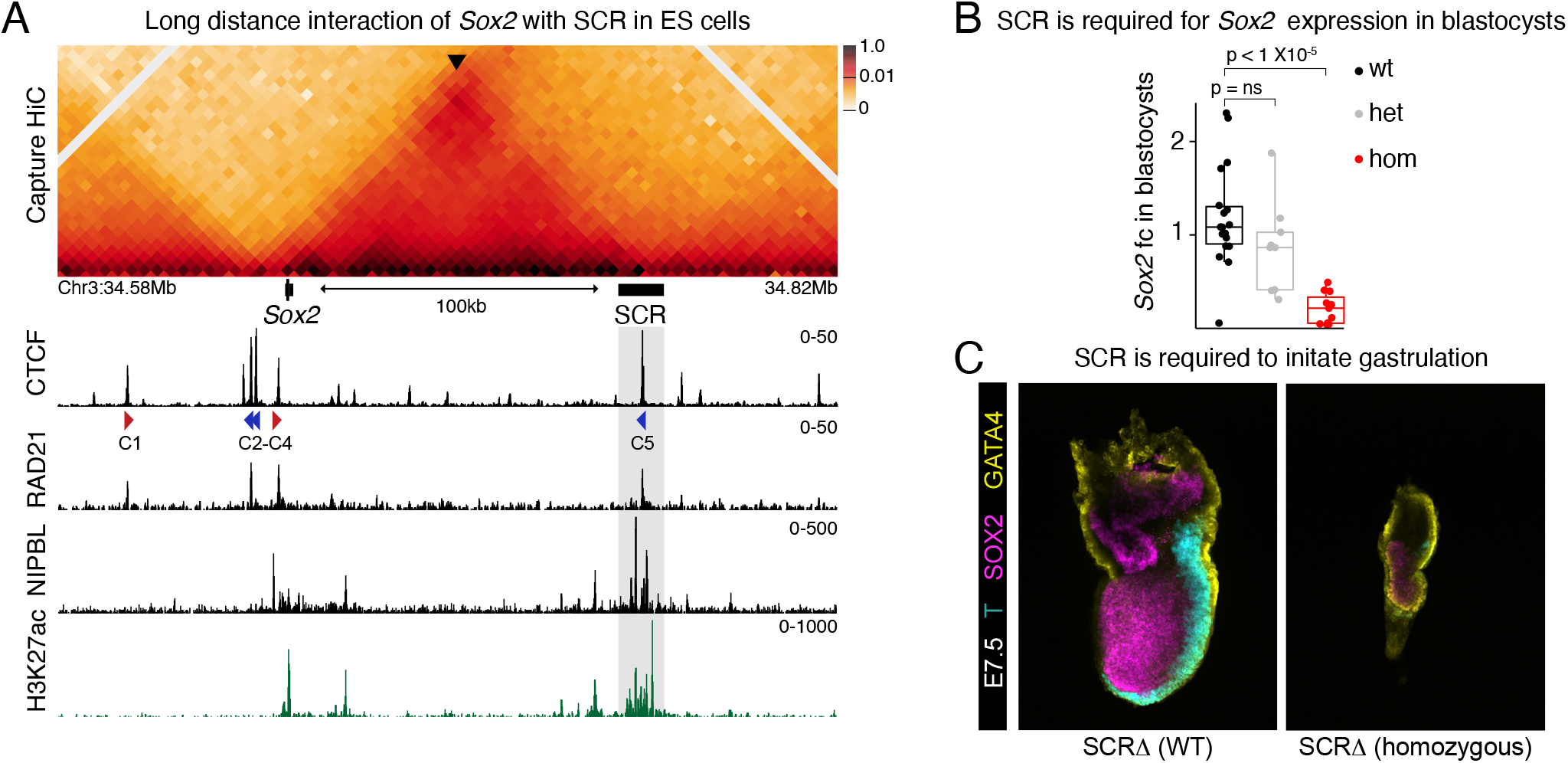
SCR is required for *Sox2* expression in epiblast cells. **A** CHiC 1D interaction frequency heatmap in wt mES cells (top). Arrowhead points to the center of the *Sox2*-SCR interaction and this corner signal overlaps with CTCF binding suggesting formation of a CTCF-mediated loop. Publicly available ChIP-seq of RAD21, CTCF, and NIPBL as well as CUT&RUN of H3K27ac in mES cells are shown below heatmap. CTCF motif orientation (red and blue arrowheads) is shown for peaks with significant CTCF motifs (pval <0.05) as detected by FIMO. Black bar and shaded box show deleted region in SCRΔ mice. **B** qPCR analysis of *Sox2* expression in blastocysts at E3.5 was done using the ΔΔCT method and Gapdh as a reference. *Sox2* expression was calculated by comparing it to the mean of all analyzed WT embryos. Each dot represents a single blastocyst. A Wilcoxon two-sided test was performed to assess statistical significance. **C** E6.5-E7.5 embryos were stained for GATA4, T, and SOX2. 8 of 9 SCRΔ homozygotes showed arrested development shortly after implantation and failed to initiate gastrulation as shown by T expression.

Here, we generated an allelic series of mice carrying targeted deletions of CTCF motifs as well as insertions of CTCF-mediated loops of varying degrees of insulation at the *Sox2* locus. We combine the phenotypic characterization of these mutant lines with Capture Hi-C (CHiC) to describe the impact of disrupting 3D chromatin structure on *Sox2* expression and its functions during development. We decouple the role of CTCF in creating insulation from its potential action in enhancer recruitment, and characterize how insulating boundaries interfere with endogenous, high-frequency enhancer-promoter contacts. The ability to overcome highly insulating borders varied between cell types and developmental processes, which allowed us to start dissecting the molecular principles determining how enhancer-promoter contacts contribute to phenotypic robustness upon perturbations of chromosomal structure.

### *Sox2* expression in pluripotent epiblast cells requires a distant enhancer known as SCR

*Sox2* is expressed in pluripotent epiblast cells before and after implantation (Avilion et al., 2003). ES cells grown in vitro resemble the pre-implantation epiblast and up to 95% of *Sox2* expression in these cells is induced by the *Sox2* Control Region (SCR). SCR is a 12 kb enhancer cluster located over 100kb downstream of the *Sox2* promoter (Li et al., 2014; Zhou et al., 2014). In ES cells, the TAD containing *Sox2* is delimited on the centromeric end by 3 CTCF motifs directly upstream of *Sox2*, and by SCR and its central CTCF motif on the telomeric end (Fig. 1A). Based on enrichment of genomic features typically associated with enhancers (ATAC-seq signal and presence of acetylated lysine 27 on histone H3 - H3K27ac), SCR is active in preimplantation epiblast pluripotent cells (Fig. S1A). Following implantation, the epiblast maintains H3K27ac enrichment at SCR but at reduced levels, which are completely lost by E11.5 in neuronal cells that express *Sox2*. Interestingly, SCR is reactivated in cells of the germ line, where pluripotency is re-established (Fig. S1A). To characterize the function of SCR in vivo during development, we injected zygotes with Cas9 ribonucleoprotein complexes containing a pair of gRNAs surrounding SCR to generate a mouse line (SCRΔ) lacking the entire enhancer cluster (Fig. S1B).

As in ES cells, homozygous loss of SCR in blastocysts at embryonic day (E) 3.5 caused a dramatic reduction of *Sox2* expression in vivo (Fig. 1B). We then asked whether SCR deletion caused most cells to lose SOX2 protein or if instead a few cells expressed normal amounts of SOX2 while others expressed none. To answer this and also assess if less SOX2 affected pre-implantation lineage fate decisions we performed a quantitative analysis of protein expression using immunofluorescence (IF) with antibodies specific for SOX2, NANOG and GATA6 (Fig. S1C) (Saiz et al., 2016). In mouse blastocysts, the first cell fate decision specifies the trophectoderm at the embryo periphery and the inner cell mass (ICM), which is characterized by high SOX2 expression. These ICM cells initially co-express NANOG and GATA6 before the exclusive expression of NANOG specifies the epiblast, and GATA6 marks cells of the primitive endoderm (Chazaud et al., 2006). Surprisingly, at E4.5 we did not find a difference in either number or localization of SOX2-expressing cells in blastocysts with homozygous loss of SCR compared to wild-type littermates. Instead, most cells showed a stark reduction in their ability to express high levels of SOX2 (Fig. S1C). In contrast to what has been described for complete SOX2 ablation (Wicklow et al., 2014), homozygous loss of SCR did not affect expression levels of NANOG and GATA6, nor the ability of embryos to specify cells with divergent expression pattern of these transcription factors (Fig. S1C).

The marked SOX2 decrease in SCRΔ homozygotes was nevertheless enough to disrupt successful implantation, and phenocopy *Sox2* knock-out animals. Embryos isolated at E6.5-E7.5 displayed developmental arrest shortly after implantation, with 8/9 homozygotes failing to express BRACHYURY (T), a marker of mesoderm differentiation/ gastrulation (Fig.1C). These data confirm that SCR is essential for expression of *Sox2* during pre- and post-implantation and establish this enhancer-promoter pair as an excellent in vivo model to study how distant enhancers achieve faithful control of gene expression during development.

### SCR regulates *Sox2* expression independently of CTCF/ cohesin-mediated loops

CTCF binds upstream of *Sox2* at a cluster of three divergent motifs (two in the negative strand and one in the positive strand) and at the center of SCR (Fig. 1A and Fig. 2A). The CTCF motif closest to *Sox2* is in convergent orientation with the motif at SCR and the interaction frequency between *Sox2* and SCR is highest at the SCR CTCF motif (see arrowhead in Fig. 1A and Fig. 2C). This suggests that CTCF-mediated loops could play an important role in this interaction and regulation of *Sox2* expression. However, deletion of the CTCF motif at SCR in ES cells cultured in vitro has no effect on *Sox2* expression (de Wit et al., 2015). It has been suggested that CTCF-mediated loops might be required to initiate interactions but clustering of similar types of chromatin can independently maintain enhancers in close contact with promoters (Alexander et al., 2019; Hsieh et al., 2021). To test whether CTCF-mediated loops are required to initiate establishment of the *Sox2*-SCR interaction during development we generated two mouse lines where CTCF motifs at the two ends of the domain were deleted. At the *Sox2* end we removed the entire cluster of three divergent CTCF motifs (named here as C2, C3 and C4). The goal of this large deletion was to ensure complete loss of cohesin retention upstream of *Sox2* and thus disrupt CTCF-mediated loops with CTCF motifs both upstream and downstream of *Sox2*. This mouse line was named CTCFΔ(C2-C4) (Fig. S2). Although this deletion removes binding sites for NANOG, OCT4, and SOX2, this region has been shown to be non-essential for mouse development (Ferri et al., 2004). The CTCF motif at SCR was replaced by a sequence of the same size and named CTCFΔ(C5) (Fig. S2).

**Figure 2:**
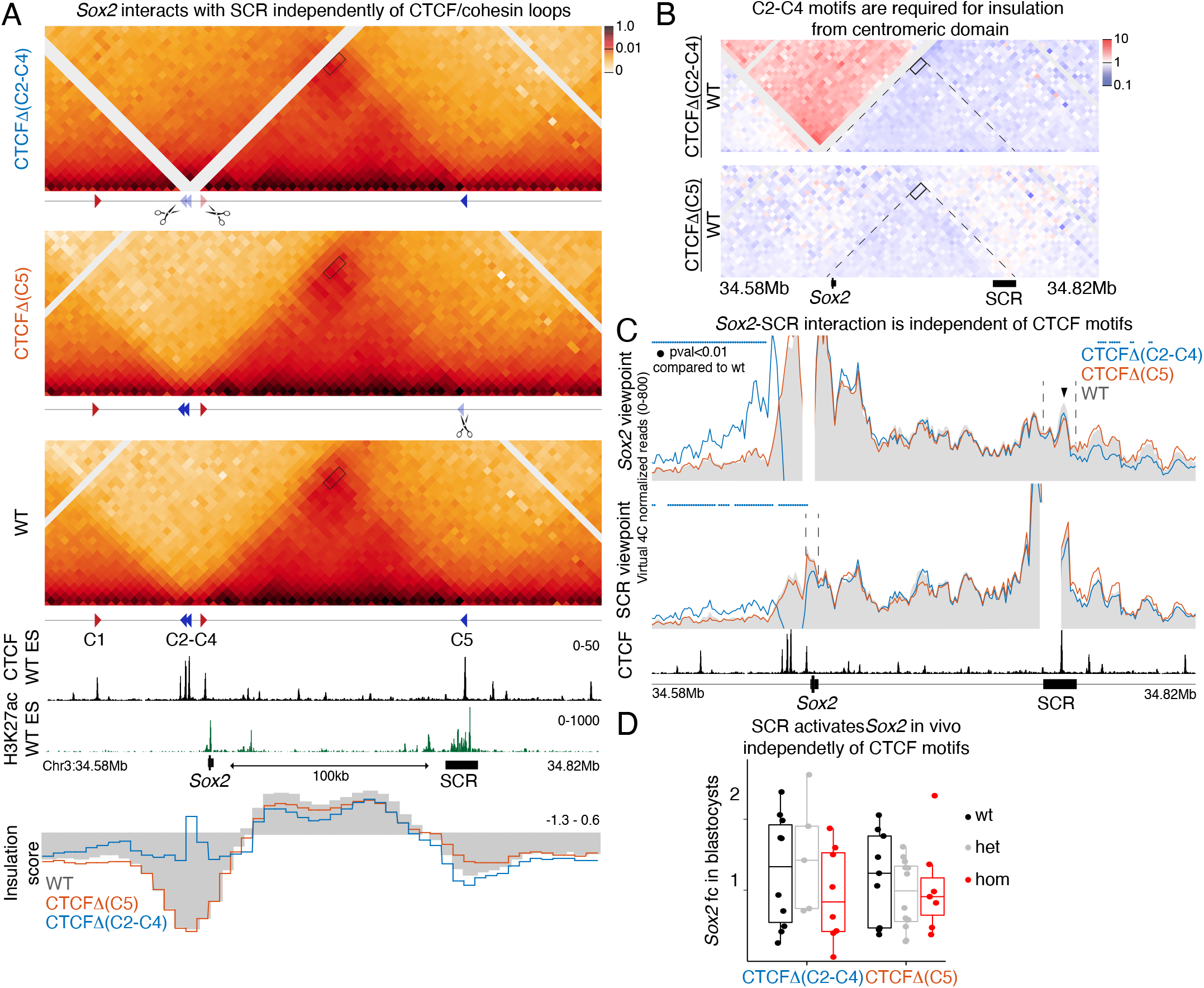
*Sox2*-SCR interaction is established independently of CTCF. **A** CHiC 1D interaction frequency heatmap in homozygotic CTCFΔ(C2-C4) and CTCFΔ(C5) mES cells compared to WT. Rectangles represent the *Sox2*-SCR interaction. Insulation scores for 5kb windows in this region are shown below publicly available CTCF and H3K27ac enrichment tracks. Lower scores represent higher insulation. **B** Differential CHiC interaction frequency heatmap. Red signal represents interactions occurring at higher frequency in mutant cell lines compared to control and blue shows interactions of lower frequency. Dotted lines represent the *Sox2*-SCR domain as detected in WT control cells. **C** Virtual 4C plot using the *Sox2* and SCR viewpoints. Region surrounding viewpoint was removed from analysis. Dotted lines highlight SCR in the *Sox2* viewpoint (top), and *Sox2* in the SCR viewpoint (bottom). Virtual 4C signal is shown as average of the 2 replicates in 5kb overlapping windows. Colored dots represent regions of statistically significant difference compared to WT (adj pval<0.01). Black arrowhead indicates region of highest intensity of the *Sox2*-SCR interaction, which overlaps with the SCR-CTCF motif. **D** qPCR analysis of *Sox2* expression in blastocysts at E3.5 was done using the ΔΔCT method and Gapdh as a reference. *Sox2* expression was calculated by comparing it to the mean of all analyzed WT embryos. Each dot represents a single blastocyst. A Wilcox-on two-sided test was performed to assess statistical significance.

To overcome low cell numbers in blastocysts, which prevent high resolution analysis of chromatin structure, we derived ES cells from homozygous blastocysts of each of these two mutant strains in parallel with wild-type littermates (Fig. 2A). For each genotype, two independently derived ES lines served as biological replicates. CHiC was then performed separately for each replicate and data pooled for visualization. Loss of CTCF motifs upstream of *Sox2* in CTCFΔ(C2-C4) caused a strong loss of insulation and fusion with the upstream domain (blue line in insulation score plot in Fig. 2A and Fig. 2B). In contrast, CTCFΔ(C5) showed a very small decrease in insulation and no noticeable change on the formation of the *Sox2*-SCR domain(orange line in insulation score plot in Fig. 2A and Fig. 2B). To facilitate quantification of changes in interaction frequencies of *Sox2* with SCR, we generated virtual 4C plots using either *Sox2* or SCR as viewpoints (Fig. 2C). This was done by plotting the frequency of all CHiC reads where one end was mapped to the area defined as viewpoint. Both viewpoints showed that loss of CTCF motifs anchoring the *Sox2*-SCR domain did not cause statistically significant changes in interaction frequency between *Sox2* and SCR. This was especially striking in CTCFΔ(C2-C4) cells where insulation with the upstream domain was completely lost but the *Sox2*-SCR interaction was not significantly affected. In both mutant lines, the highest *Sox2*-SCR interaction signal was still centered on the CTCF motif (square in Fig. 2A and arrowhead in Fig. 2C). This region of maximum signal appeared slightly reduced compared to WT cells but did not meet our threshold for significance suggesting that the CTCF motif itself is not the cause of higher interaction frequency.

In line with high *Sox2*-SCR interactions in the absence of CTCF motifs, we did not detect statistically significant differences (pval<0.05) in *Sox2* expression in blastocysts from either CTCFΔ(C2-C4) or CTCFΔ(C5) mutants (Fig. 2D). Together, these data show that CTCF-mediated loops are not only dispensable to maintain *Sox2* contacts with SCR in vitro but are also not required for de novo establishment of this interaction, or for *Sox2* expression in vivo. The very slight decrease in interaction frequency over the CTCF motif at SCR also supports that CTCF-mediated loops play only a limited role in establishment of the Sox2-SCR interaction. Our data show that generation of insulation by CTCF-mediated loops can be decoupled from their role in establishment of interactions between enhancers and promoters even when these regulatory elements are found directly adjacent to domain boundaries.

### High affinity *Sox2*-SCR interaction can bypass CTCF-mediated loops

Domain boundary deletions that lead to formation of new regulatory contacts and transcriptional activation suggest that CTCF/cohesin-mediated loops can restrict the range of enhancer action (Dowen et al., 2014; Hanssen et al., 2017; Lupianez et al., 2015; Narendra et al., 2015). However, it is less clear whether such loops can disrupt native enhancer-promoter interactions that are responsible for tissue specific gene expression during development (Despang et al., 2019; Rodriguez-Carballo et al., 2020). To address this, we perturbed the Sox2 locus by inserting CTCF motifs in combinations that we predicted would lead to loops of different insulation strength. We targeted a cassette of three CTCF motifs to two locations between Sox2 and SCR (Fig. S3A). We assumed these sites to not contain regulatory elements based on conservation and analysis of ENCODE datasets. The CTCF cassette consisted of tissue-invariant, strongly-bound CTCF motifs, that retain cohesin efficiently (Redolfi et al., 2019). The mouse line CTCFi3x(-) was generated by targeting the cassette to contain CTCF motifs on the negative strand. This should create a loop with the CTCF motif in the positive strand, upstream of Sox2 (Fig. 3A and S3A). In contrast, the CTCFi3x(+) line was created by inserting the cassette closer to SCR with CTCF motifs on the positive strand to form a loop with the CTCF motif at SCR. Enrichment of the cohesin loader NIPBL suggests that cohesin can be loaded at both Sox2 and SCR (Fig. 1A). Therefore, the two insertions were expected to bind CTCF in the correct orientation to halt cohesin extrusion (Vos et al., 2021). We also generated the CTCFi3x(-);3x(+) line by combining both insertions, predicting the formation of two ectopic loops. Finally, CTCFi18x(+) mice carrying 6 copies of the transgene for a total of 18 motifs, were used to assess the ability of CTCF binding at high density to block cohesin extrusion.

**Figure 3:**
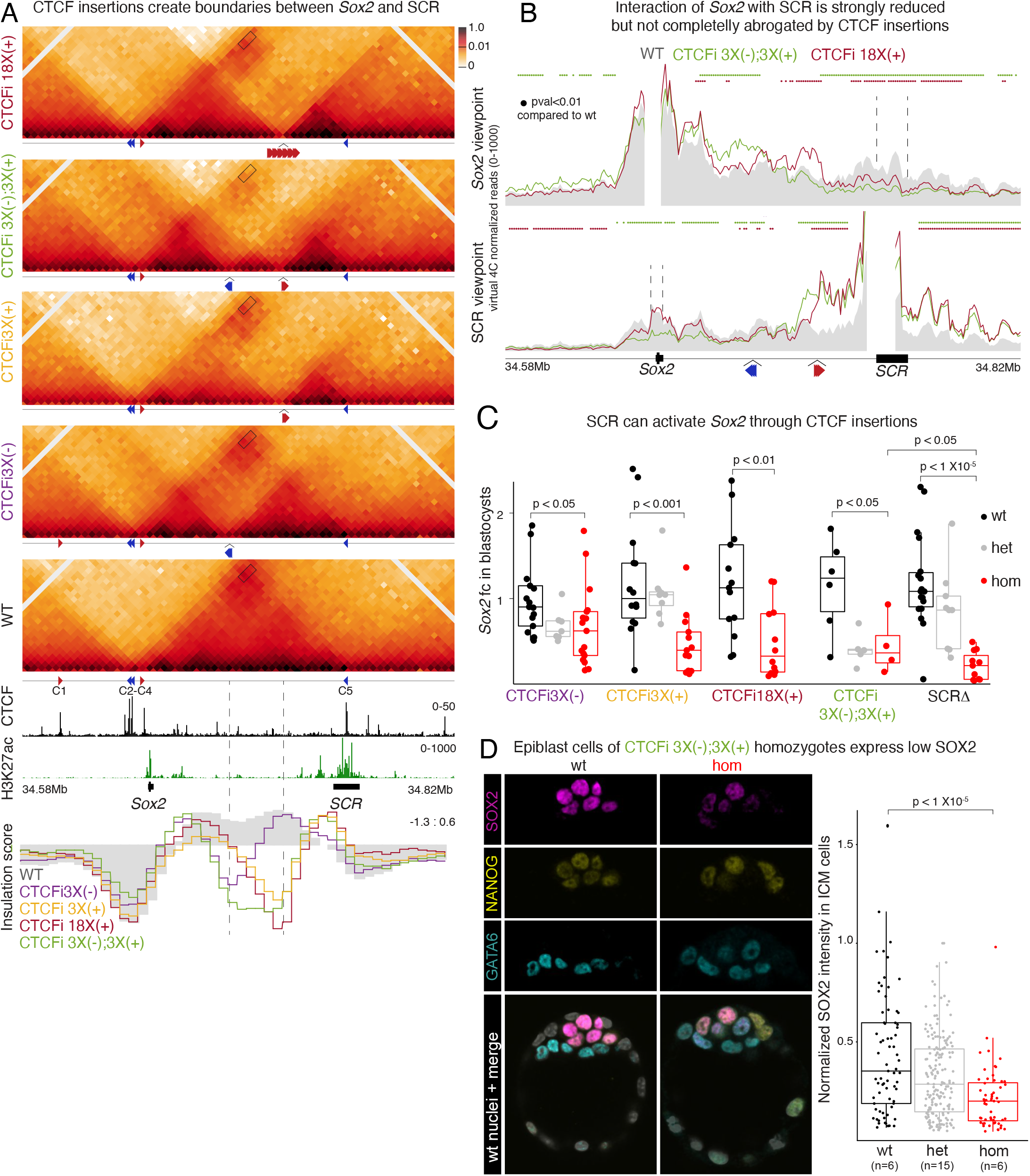
SCR can bypass strong CTCF-mediated loops to induce *Sox2*. **A** CHiC 1D interaction frequency heatmap in homozygotic mES cells of the CTCFi3x(-), CTCFi3x(+), CTCFi18x(+), and CTCFi3x(-);3x(+) strains compared to WT. Rectangles show the *Sox2*-SCR interaction. Inserted CTCF motif orientation and position in each mutant is shown below the plots. Insulation scores for 5kb windows shown below publicly available CTCF and H3K27ac tracks. Dotted lines show CTCF insertion sites. **B** Virtual 4C plot using the *Sox2* and SCR viewpoints. Region surrounding viewpoint was removed from analysis. Virtual 4C signal is shown as average of the 2 replicates in 5kb overlapping windows. Colored dots represent regions of statistically significant difference compared to WT (adj pval<0.01). **C** qPCR analysis of *Sox2* expression in blastocysts at E3.5 was done using the ΔΔCT method and Gapdh as a reference. Sox2 expression was compared to the mean of all WT embryos. Each dot is a blastocyst and a Wilcoxon two-sided test assessed statistical significance. **D** IF of blastocysts with antibodies targeting GATA6, NANOG and SOX2. Each dot represents the signal intensity of a single cell normalized by the cell with highest intensity in heterozygotes. Number of embryos analyzed is shown below the plot. A Wilcoxon two-sided test was performed to assess statistical significance.

To determine the effect of ectopic loop formation on Sox2 expression and enhancer-promoter interactions, we performed CHiC on ES cells derived from homozygous blastocysts of each line. These data showed that our predictions based on the loop extrusion model were correct. Insertion of CTCF motifs in the CTCFi3x(-), CTCFi3x(+), and CTCFi18x(+) mouse lines resulted in the formation of two subdomains between Sox2 and SCR, with a new boundary at the location of transgene insertion (Fig.3A). Similarly, the double CTCF cassette in CTCFi3x(-);3x(+) mice led to the formation of 3 subdomains. Based on insulation score, the greatest local increase in insulation was seen in CTCFi18x(+) cells, while CTCFi3x(-);3x(+) cells showed a larger, highly insulating boundary (Fig. 3A and S3B). Surprisingly, despite formation of a strong boundary in CTCFi3x(-) and CTCFi3x(+) cells, the interaction of Sox2 with SCR was only minimally affected (Fig. 3B and S3C). Sox2-SCR contacts were more perturbed in CTCFi18x(+) cells and even further in cells where CTCF motifs were inserted across a larger region in divergent orientation in the CTCFi3x(-);3x(+) line. Despite this decrease, in CTCFi3x(-);3x(+) cells we could still detect the Sox2-SCR interaction, which showed higher CHiC signal between the promoter and the enhancer cluster compared to adjacent regions (Fig. 3B and S3C).

We then assessed how the changes in 3D structure and reduced Sox2-SCR interactions affected Sox2 expression in blastocysts. Although homozygous embryos of all lines had significantly (pval<0.05) reduced Sox2 expression, none of our mutants showed a loss comparable to what we quantified upon SCR deletion (Fig. 3C). Importantly, the two lines that showed the strongest reduction in interactions, CTCFi18x(+) and CTCFi3x(-);3x(+), also showed the greatest reduction in Sox2 expression. We hypothesized that Sox2 expression was higher in blastocysts with CTCF insertions than in those without SCR because the dynamic nature of CTCF binding—and consequently loop formation—would allow sporadic SCR-Sox2 contacts leading to stronger expression in a few cells. To test this, we again quantified single-cell protein expression in blastocysts, focusing on CTCFi3x(-);3x(+) mutants as they showed the strongest decrease in SCR contacts and Sox2 expression. Similar to the SCRΔ line, CTCFi3x(-);3x(+) homozygous blastocysts showed no defect in specification of epiblast and primitive endoderm cells or in the total number of SOX2-expressing cells (Fig. 3D). Instead, we found that all epiblast cells expressed less SOX2 suggesting that the CTCF motif insertion homogenously reduced SCR contacts with Sox2 and its transcription. In contrast to the post implantation lethality of SCRΔ, we found that CTCFi3x(-);3x(+) animals, as well as homozygotes of all other lines, completed gastrulation, and initiated organogenesis without any overt phenotypes (see Fig. 4D for examples of mid-gestation E9.5-10.5 embryos).

**Figure 4:**
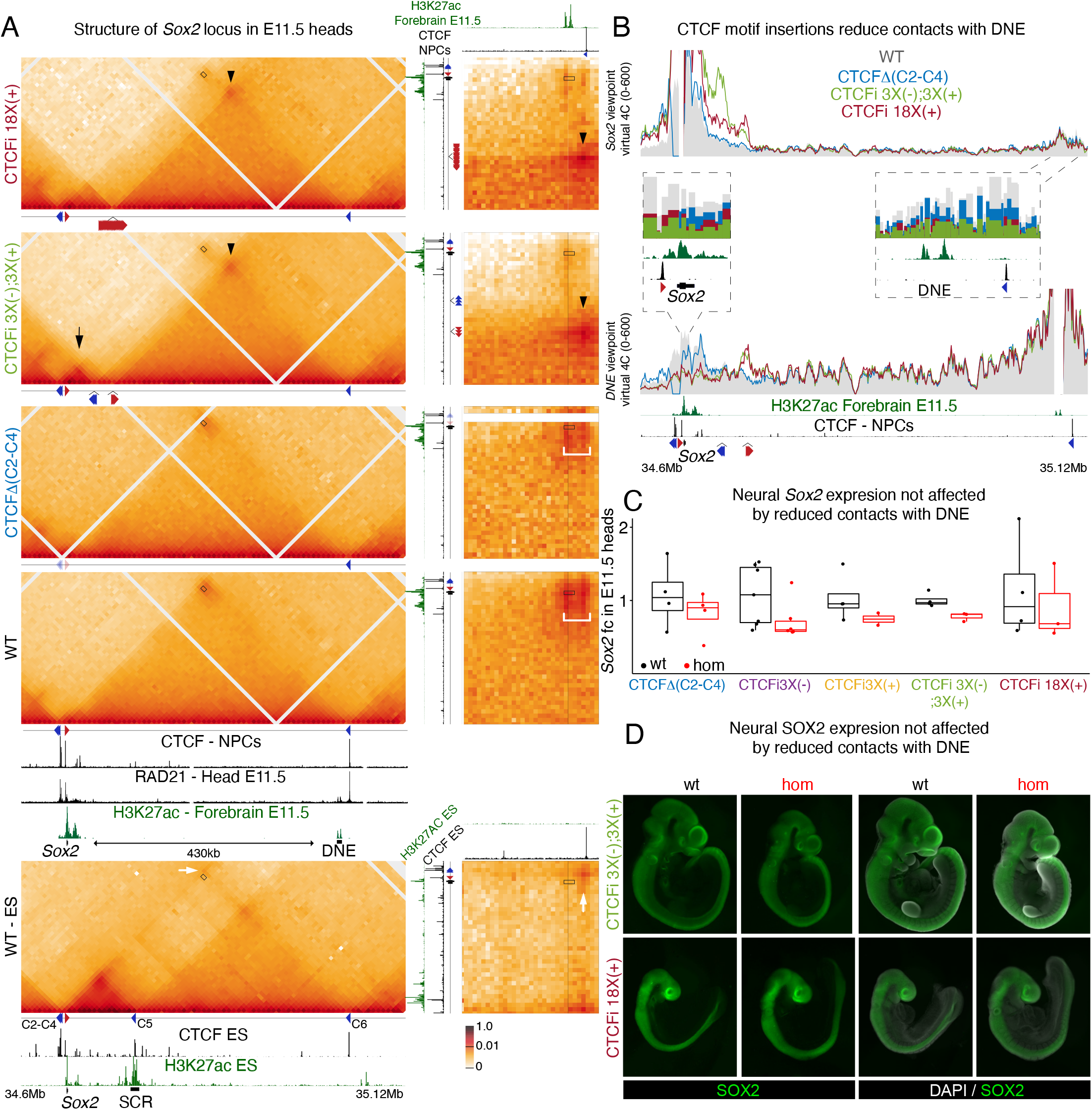
Interaction of *Sox2* with long-distance neural enhancers also bypasses CTCF-mediated insulation. **A** CHiC 1D interaction frequency heatmap in E11.5 heads. Data of WT ES cells is shown for comparison (bottom). Insets on the right show 2D interaction heatmaps highlighting interactions between regions surrounding *Sox2* and DNE. CTCF ChIP-seq publicly available data was obtained from in vitro differentiated neural progenitor cells. RAD21 ChIP-seq was performed on E11.5 heads isolated as for CHiC. 2D insets show the same tracks in x and y axis but at different locations. Rectangles represent the *Sox2*-DNE interaction, arrowheads represent loops with CTCF downstream of DNE established by transgene insertions, black arrow represents loops with CTCF upstream of *Sox2* established by transgene insertions, white arrow represents loops in ES cells between CTCF upstream of *Sox2* and CTCF downstream of DNE, white brackets in the insets highlight the region between DNE and CTCF. **B** Virtual 4C plot using Sox2 and DNE viewpoints using 5kb overlapping windows and signal is shown as average of the 2 replicates of each genotype. Region surrounding viewpoint was removed from analysis. Insets show signal at DpnII fragments. **C** qPCR analysis of *Sox2* expression in E11.5 heads was done using the ΔΔCT method and Gapdh as a reference. *Sox2* expression was compared to the mean of all WT embryos. Each dot is one embryo and a Wilcoxon two-sided test assessed statistical significance. **D** IF of E9.5-10.5 embryos stained with an antibody targeting SOX2

In sum, these data show that SCR regulates Sox2 through a high affinity interaction that can bypass strong insulating CTCF-mediated loops. The impact on Sox2 expression correlated with the ability of CTCF insertions to disrupt this interaction highlighting that proximity of enhancers to promoters can play a determinant role in expression levels. Furthermore, our data show that enhancer-promoter affinity contributes to phenotypic robustness as the vastly reduced SCR contacts still induced enough SOX2 expression for successful implantation. These data also show that a high-density cluster of CTCF motifs in the same orientation, as in CTCFi18x(+) mice, can lead to stronger local insulation but a combination of CTCF motifs in divergent orientations, such as in CTCFi3x(-);3x(+), blocks enhancer-promoter contacts more efficiently by forming loops with both upstream and downstream CTCF motifs.

### CTCF-independent interactions with long-distance neural enhancers also bypass CTCF boundaries

*Sox2* is one of the earliest neural markers and is essential for successful neurogenesis (Favaro et al., 2009; Mercurio et al., 2021). To describe the chromatin structure of the *Sox2* locus in cells that adopt a neural fate we performed CHiC on tissues isolated from the heads of E11.5 embryos in which forebrain, midbrain, and hindbrain were pooled. This showed that the 3D structure of *Sox2* in neural cells is very different from the epiblast. Along with loss of SCR activity, the CTCF motif at SCR becomes unoccupied, cohesin is no longer retained and the strong interaction domain is lost (Fig. 4A and S4A). In neural tissues, *Sox2* forms a larger (∼430 kb) domain that is delimited at its telomeric end by a CTCF motif in the negative strand and a region that shows neural-specific H3K27ac enrichment that has been suggested to regulate *Sox2* expression in these cells (Beagan et al., 2017; Bonev et al., 2017). For simplicity we refer to this region as distal neural enhancer (DNE). Proximal to the *Sox2* locus, several neural enhancers have been identified (Uchikawa and Kondoh, 2016) that also show specific activity in neuronal cells (Fig. S4A). Like in ES cells, CHiC data alone is inconclusive to determine if CTCF-mediated loops play an essential role in the *Sox2*–DNE interaction as the signal at the domain corner encompasses both DNE and the CTCF peak (bracket in inset of WT heads in Fig. 4A). The inactive state of DNE in ES cells, reveals a clear corner signal between the CTCF motifs upstream of *Sox2* and those downstream of DNE (white arrow in ES cells of Fig. 4A), which supports that both CTCF peaks can retain cohesin and form a large loop.

We then asked whether the CTCF motifs upstream of *Sox2* function as anchors to facilitate contacts with proximal enhancers and DNE. Additionally, we assessed whether DNE, similarly to SCR, could bypass CTCF-mediated loops to interact with *Sox2* in our CTCF insertion mouse lines. For this, we did CHiC with chromatin isolated from the head of individual E11.5 embryos, using two replicates of each genotype. As in ES cells, deletion of the CTCF motifs upstream of *Sox2*, in CTCFΔ(C2-C4) homozygous embryos caused loss of insulation and fusion with the upstream domain (Fig. 4A, S4B and S4C). While contacts with the CTCF motif downstream of DNE were reduced to approximately half (inset in Fig. 4B in the *Sox2* viewpoint), the interaction with DNE was only minimally affected (bracket in insets of Fig. 4A and Fig. 4B), suggesting that CTCF is also not essential for the interaction in neural cells.

CTCF insertions generated new loops between the CTCF motif downstream of DNE in the negative strand, and the motifs targeted to the positive orientation in CTCFi3x(+), CTCF18x(+), and CTCFi3x(-);3x(+) embryos (arrowheads in Fig. 4A and S4A). Similar to ES cells, homozygotes of the CTCF18x(+) line showed the strongest insulation score at the integration site, and CTCFi3x(-);3x(+) generated a larger, highly insulating boundary (Fig. S4B and S4C). Interestingly, interactions with proximal neural enhancers increased in CTCFi3x(-) and CTCFi3x(-);3x(+) embryos where CTCF was targeted in the negative strand because of formation of a strong interacting domain with CTCF upstream of Sox2 (black arrows in Fig. 4A, S4A and S4C). As in ES cells, upon introduction of CTCF-mediated loops, Sox2 and DNE interacted less frequently with the starkest decrease seen in CTCFi3x(-);3x(+) homozygotes (Fig. 4A, 4B and S4B). However, even in these embryos, Sox2-DNE contacts were not abrogated completely.

Measuring Sox2 expression in the same tissues used for CHiC revealed that Sox2 expression was not significantly affected (pval<0.05) either by the loss of CTCF binding upstream of its promoter or by the introduction of ectopic loops (Fig. 4C). In addition, whole mount IF with an anti-SOX2 antibody confirmed that SOX2 expression in neural tissues remained largely unaltered (Fig. 4D and S4E). Unperturbed Sox2 expression upon loss of the telomeric anchor in CTCFΔ(C2-C4) mutants and the increase in short range contacts in CTCFi3X(-) and CTCFi3x(-);3x(+) homozygotes, suggest that CTCF loops do not play a role in regulation of Sox2 and may reflect even higher resilience to structural perturbations in neural cells than in the epiblast (Fig. 4C). The absence of significant changes in Sox2 expression despite the stark reduction in contacts with DNE, could also be explained by compensatory roles by Sox2 proximal enhancers. It is also possible that DNE itself has a negligible impact on Sox2 expression despite their strong interaction in WT embryos and the tissue specific activity of this putative enhancer. Nonetheless, just like in ES cells, Sox2-DNE contacts are not abrogated completely by introduction of ectopic CTCF-mediated loops.

### *Sox2* regulation by distal enhancers in the anterior foregut is more sensitive to structural perturbations

Despite closer inspection of CTCFi18X(+) and CTCFi3x(-);3x(+) IFs, we noticed that insertion of CTCF motifs caused SOX2 to become undetectable in the anterior foregut (AFG) at E9.5 (white bracket in Fig. 5A). At this stage the AFG is an epithelial layer located between the developing heart and the neural tube and is marked by strong *Sox2* expression. At E10.5 the AFG separates, and the most ventral tube will form the trachea and lungs, while dorsally it gives rise to the esophagus and stomach (Edwards et al., 2021). Mice with hypomorphic *Sox2* alleles and patients with heterozygous SOX2 mutations fail to separate the AFG and develop a tracheoesophageal fistula where a single tube connects the pharynx to the lungs and stomach (Que et al., 2007; Teramoto et al., 2020; Zenteno et al., 2006). Consistent with the disruption of SOX2 expression in the AFG, at E13.5 CTCFi3x(-);3x(+) homozygotes developed a tracheoesophageal fistula and did not express SOX2 in any AFG-derived tissues (Fig. 5B and Fig. S5B). NKX2.1, a transcription factor that is normally expressed only in respiratory organs (trachea and lungs), was found across the entire fistula including the stomach (Fig. 5B). Failure in separating the trachea and esophagus was followed by perinatal lethality of CTCFi3x(-);3x(+) homozygotes. Although the three possible genotypes were recovered at the expected ratio at E18.5, all homozygous pups were found dead within a few hours of being born (Fig. S5A). In line with absence of SOX2 expression at E9.5, we only recovered three CTCFi18X(+) homozygous pups alive at P0 (out of 15 expected) and none at weaning. Importantly, loss of CTCF motifs upstream of *Sox2* in CTCFΔ(C2-C4) homozygotes did not affect *Sox2* AFG expression (Fig. S4E), indicating that also in these tissues, *Sox2* can be induced independently of CTCF-mediated loops adjacent to its promoter.

**Figure 5:**
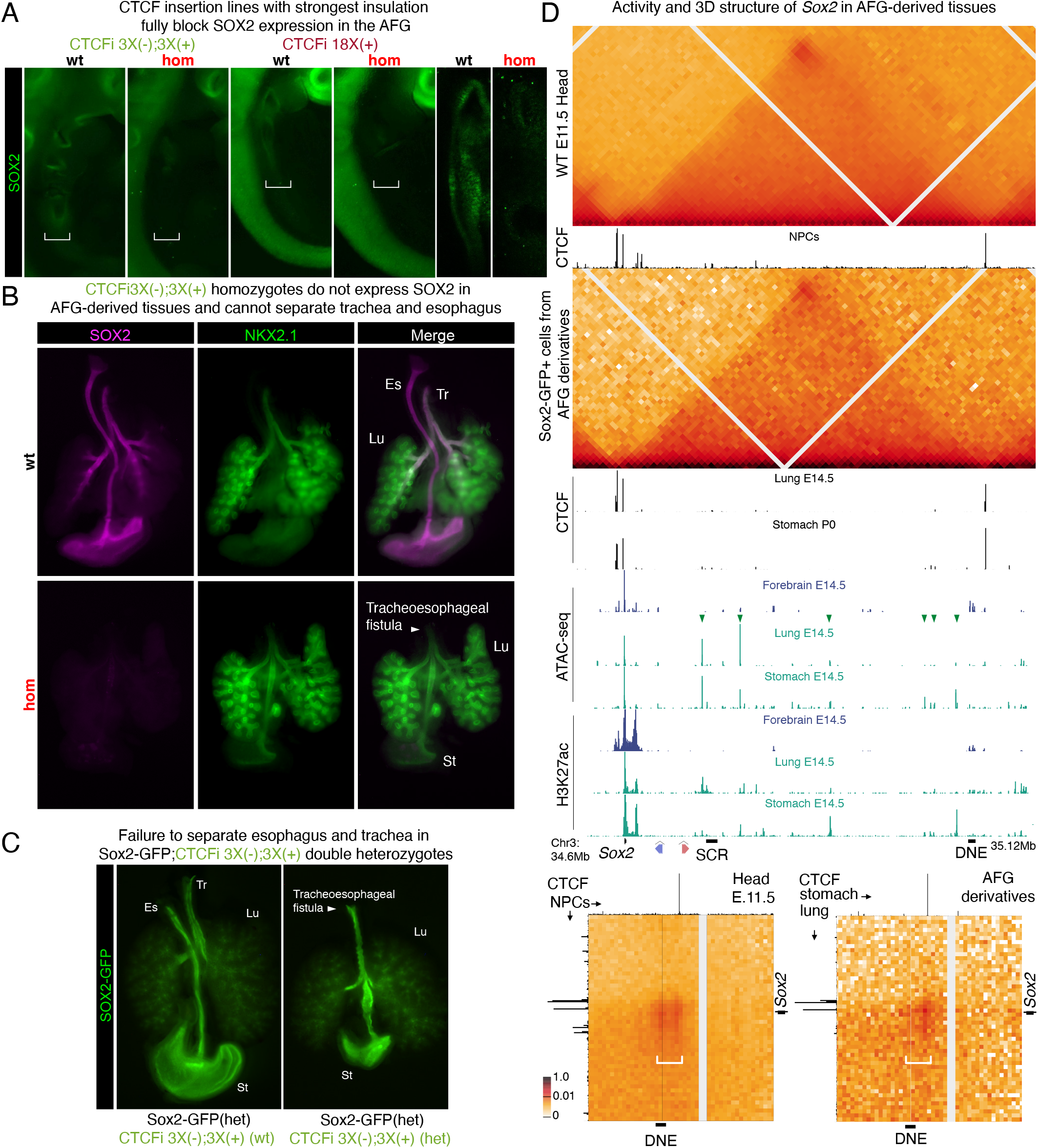
CTCF-loops can completely insulate *Sox2* from its AFG-specific enhancers. **A** IF of E11.5 embryos stained with an antibody targeting SOX2. Bracket highlights the AFG. First 4 images were taken with a dissection microscope and the two highlights on the right with a confocal microscope. **B** IF with antibodies targeting SOX2 and NKX2.1 using dissected E13.5 AFG-derived tissues. Tr-trachea Es-esophagus Lu-Lungs St-stomach. **C** GFP expression in AFG-derived organs dissected from E15.5 fetuses originating from crosses between CTCFi3x(-);3x(+) and *Sox2-GFP* heterozygous mice. **D** CHiC 1D interaction frequency heatmap in WT E11.5 heads (top) and GFP+ cells from E11.5-12.5 AFG-derived tissues dissected from *Sox2-GF*P heterozygotes (bottom). Publicly available CTCF, ATAC-seq and H3K27ac enrichment data from different tissues is shown below heatmap. Green arrowheads indicate putative regulatory elements with tissue-specific activity in AFG-derivatives. Insets on the right show a 2D interaction heatmap where Y axis shows region centered around *Sox2* and x axis a region around a CTCF motif downstream of DNE. For the E11.5 head inset, CTCF data from NPCs is shown in both axis. For AFG-derived tissues, CTCF from stomach is shown in x axis and from lungs in the y axis.

The striking AFG phenotype seen in CTCFi3x(-);3x(+) and CTCFi18X(+) homozygotes, prompted us to examine viability of all homozygotes to better understand how the structural mutations represented in our allelic series impacted animal development. We recovered CTCFΔ(C2-C4), CTCFΔ(C5), CTCFi3X(-), and CTCFi3(+) homozygotes at weaning without significant (pval<0.01) deviation from expected ratio indicating they are compatible with animal development (Fig. S5A). The viability of homozygous CTCFi3x(-) and CTCFi3x(+) animals provides strong evidence that insertion of CTCF cassettes did not disrupt regulatory elements and that the phenotypes seen in CTCFi3x(-);3x(+) and CTCFi18X(+) homozygotes are caused by perturbations to the chromatin structure of the *Sox2* locus. In agreement with our observation that 1 of 9 post-implantation E6.5 mutant embryos initiated gastrulation (Fig. 1C), we recovered a few SCRΔ homozygotes at weaning but at a highly reduced frequency compared to the expected ratio (Fig. S5A). This could be explained by SCR losing activity following implantation (Fig. S1A), suggesting that embryos that successfully implant despite SCR deletion can complete development. The fully penetrant perinatal lethality of CTCFi3x(-);3x(+) and CTCFi18X(+) animals is consistent with the AFG fusion phenotype occurring with complete penetrance in these strains. We also found craniofacial phenotypes, such as cleft palate, in some CTCFi3x(-);3x(+) and CTCFi18X(+) homozygotes at E18.5/P0 (Fig. S1C). As SOX2 has also been implicated in craniofacial development (Langer et al., 2014; Mandalos et al., 2014) this points to yet other specific cell types where the impact of highly insulating boundaries is more severe. To assess if a further decrease of *Sox2* levels would lead to a phenotype during implantation, we set up crosses between heterozygotes of the CTCFi3x(-);3x(+) and Sox2-GFP lines (Arnold et al., 2011). The latter carry an allele where the *Sox2* coding sequence was substituted by GFP. As such, double heterozygous embryos have only one functional copy of *Sox2*, which also contains the double CTCF cassette insertion. Surprisingly, embryos carrying both alleles (Sox2-GFP in one chromosome 3 and CTCFi3x(-);3x(+) in the other) successfully implanted and died shortly after birth. As expected, fetuses of these genotypes developed tracheoesophageal fistula (Fig. 5C). These results provide further evidence of the extreme resilience of blastocysts to low *Sox2* levels in contrast to the higher susceptibility during AFG development.

We then focused on understanding why CTCF insertions had a stronger impact on *Sox2* expression that led to increased phenotypic susceptibility in the AFG compared to pre-implantation development and neurogenesis. We hypothesized that tissues with more *Sox2* expression might be more resilient to loss of contacts with distal enhancers. It is hard to compare gene expression in blastocysts to later stages due to the wide differences in cell numbers that may affect precise quantification. However, single-cell RNA-seq at midgestation revealed that *Sox2* is less expressed in the AFG than in neural cells (Fig. S5D) (Ibarra-Soria et al., 2018). AFG-specific regulatory elements also showed different characteristics compared to epiblast and cells of the neural fate. In lung and stomach of E14.5 embryos, we found several regions downstream of our CTCF insertion locations with AFG-specific enrichment of H3K27ac ChIP-seq and ATAC-seq signal compared to the forebrain where *Sox2* is also expressed (green arrowheads in Fig 5D). Unlike in ES cells where SCR displays high density of marks associated with regulatory activity, putative AFG-specific enhancers were spread through a much larger domain. The most proximal AFG-specific peak was found ∼100kb away from *Sox2*, close to SCR. The most distal peak was located 400kb away, just upstream of DNE. ATAC-seq and H3K27ac were also enriched more proximally to *Sox2*, with a similar pattern to what is detected in neural tissues suggesting that these enhancers may work in a non-tissue specific manner. We also characterized the 3D structure of the *Sox2* locus in AFG-derived tissues. CTCF binding in the developing stomach and lungs is remarkably similar to forebrain, with binding just upstream of *Sox2* and downstream of DNE without visible peaks in between (Fig. 5D). To define interaction frequencies, we manually dissected AFG-derived tissues from E11.5-E12.5 Sox2-GFP heterozygous embryos and then purified the GFP+ population of *Sox2*-expressing cells to perform CHiC (Fig. 5D). We were especially careful to ensure that no neural tissue was included in the manual dissection. Although CHiC coverage was lower than in ES cells and neural tissues because of fewer SOX2-GFP expressing cells, our data revealed a highly interacting domain clearly delimited by the CTCF peaks in lung and stomach primordia. Our data also points to *Sox2* interacting across the entire domain at high frequency rather than specific interactions with putative AFG enhancers (Fig. 5D). Unlike neural cells, where DNE is active and the domain corner signal spans the distal CTCF motif and DNE, in AFG-derived tissues we detected increased interaction frequency specifically between the CTCF binding sites (white brackets in insets in Fig. 5D). As discussed in more detail below, we propose that baseline transcriptional levels and enhancer density may influence the ability of distal enhancers to overcome physical barriers and maintain faithful gene development..

## DISCUSSION

The physical distance between enhancers and promoters can influence transcriptional output, the folding of the genome can impact gene expression (Beagan et al., 2020; Deng et al., 2014; Rinzema et al., 2021; Symmons et al., 2016; Zuin et al., 2021). With cohesin, CTCF mediates the formation of DNA loops and has been proposed to both facilitate or restrict enhancer-promoter interactions. Yet, how CTCF-mediated loops influence such regulatory interactions and consequently transcription is not fully understood and remains a topic of intense research. *Sox2* has often been used as an in vitro model to examine the impact of chromatin structure on gene expression (Alexander et al., 2019; Huang et al., 2021). However, it wasn’t clear how the different forces of nuclear organization influence *Sox2* expression and its many physiological roles. Here, we addressed this question by developing an allelic series of mouse mutants carrying deletions of CTCF binding sites or targeted insertions of CTCF motifs at the Sox2 locus. We dissected the impact of CTCF-loops on chromatin structure, *Sox2* expression, and across different developmental processes. Our work decouples generation of insulation by CTCF-mediated loops from a role in enhancer recruitment, describes an unanticipated high-affinity of enhancers towards its target promoters, and starts dissecting the molecular characteristics that determine resilience of regulatory interactions to ensure faithful gene expression.

The description of TADs and the frequent presence of CTCF motifs at their borders have suggested a simple model that divides the influence of CTCF-mediated loops on gene expression into two main forces: insulation that restricts the range of enhancer-promoter interactions, and promotion of physical contacts between these two types of regulatory elements (Merkenschlager and Nora, 2016; Sexton and Cavalli, 2015). Our data show that although CTCF-mediated loops form domains that insulate *Sox2* from interactions with neighboring regions, these loops are not required for *Sox2* to interact with its distal tissue-specific enhancers. In addition, *Sox2* expression was not affected by loss of CTCF-mediated loops at any of the developmental stages we assessed. These observations decouple enhancer-promoter contacts from CTCF-mediated insulation even when genes are located close to CTCF motifs that function as loop anchors, and suggest that CTCF may have a limited role in promoting enhancer recruitment. This agrees with other loci-specific dissections of TAD borders and CTCF-mediated loops, where boundary loss led to ectopic enhancer-promoter contacts, induced gene expression, and caused developmental phenotypes while disruption of loops anchoring enhancer-promoter interactions had much milder effects (Dowen et al., 2014; Flavahan et al., 2016; Laugsch et al., 2019; Narendra et al., 2015; van Bemmel et al., 2019; Williamson et al., 2019). In line with these observations, in vitro acute depletion of proteins involved in CTCF-mediated loop extrusion also does not lower transcription rates (Hsieh et al., 2021; Nora et al., 2017; Rao et al., 2017). Perhaps more surprisingly, these studies also failed to detect vast upregulation of gene expression as may be expected upon boundary loss and potential exposure of genes to ectopic enhancers. The contrast with more striking effects when insulation is lost in vivo, could be related to higher stability and homogeneity of enhancer-promoter contacts of cells cultured in vitro in a steady state condition. It will be important to characterize how such acute depletions may affect gene expression via insulation loss, in cells undergoing differentiation processes. Our data revealed a surprisingly high affinity of *Sox2* towards its tissue-specific enhancers. For example, insertion of CTCF motifs between *Sox2* and SCR formed strong insulating boundaries, reduced enhancer-promoter contacts, and decreased transcriptional output. However, these boundaries did not completely abolish Sox2-SCR interactions or enhancer-driven transcriptional activation. Indeed, some enhancer-promoter contacts were able to overcome insulating boundaries in embryos carrying a high density of CTCF motifs in the same orientation as in the CTCFi18X(+) line. Importantly, contacts were more strongly disrupted in a mouse line with fewer motifs but positioned in opposing orientations. This finding supports the hypothesis that the clustering of divergent CTCF motifs often observed at TAD borders is an essential feature of nuclear organization to delimit the range of enhancer action (Anania et al., 2021; Chang et al., 2021).

Our results point to locus and tissue-specific susceptibility to perturbations of chromatin structure. In blastocysts, our strongest ectopic insulation construct did not fully block SCR from interacting with *Sox2*, and embryos expressed enough SOX2 to complete implantation. In contrast, in the AFG, the same mouse lines fully phenocopied loss of function *Sox2* alleles and no SOX2 protein could be detected. We propose two reasons that could explain the tissue-specific differences in transcriptional insulation and phenotypic robustness to perturbation of chromatin structure. On one hand, expression levels could play an important role in determining how much loss of enhancer-promoter contacts affects physiological outcomes. As the impact of enhancers on transcription is not linear, higher transcriptional levels may contribute to ensure a minimal transcriptional output upon a reduction of contact frequencies. On the other hand, we find that regions with AFG tissue-specific activity are spread across 400kb, in contrast to the SCR where genomic regions with high transcriptional-activity are clustered in a small genomic area (12kb). Enhancer clustering likely increases local concentration of transcription factors, co-regulatory complexes, and chromatin with similar modifications (Hnisz et al., 2017; Oudelaar et al., 2018). This in turn may facilitate and strengthen the formation of physical hubs with target promoters. We therefore suggest that clustering of regulatory elements contributes to the strength of enhancer-promoter contacts which in turn contributes to the ability to overcome barriers such as CTCF-mediated insulation.

Finally, high affinity interactions between enhancers and promoters such as those we report here may have provided a selective advantage during evolution by ensuring faithful tissue-specific expression in the face of structural perturbations that interfere with regulatory contacts. Such perturbations can include chromosomal abnormalities such as amplifications or inversions but also acquisition of CTCF motifs by transposition of repeat elements. For example, SINE B2 transposons can contribute to new CTCF binding sites and could endanger efficient regulatory contacts following transposition (Choudhary et al., 2020; Schmidt et al., 2012). Ensuring maintenance of endogenous pre-established contacts could provide additional flexibility for perturbations to be either positive or negatively selected, depending on physiological outcomes. In fact, an extreme example in drosophila has shown that dramatic chromosomal rearrangements do not necessarily lead to massive transcriptional dysregulation (Ghavi-Helm et al., 2019). An important challenge going forward will be to better define the proteins, complexes and CTCF-independent mechanisms that establish such specific high-affinity regulatory interactions. Our study argues against models that rely on chromosome tracking in cis between enhancers and promoters (Furlong and Levine, 2018). Instead, our data support looping or other mechanisms that promote regulatory contacts and that may, at least partially, overcome barriers created by CTCF and cohesin. Based on our observations, it is likely that such mechanisms are tissue-specific or vary with the type and strength of enhancers. The Sox2 locus and its regulation by different long-distance enhancers could prove to be a very useful model for characterization of such processes both in vivo, and in vitro through genome-wide loss-of-function screens.

## METHODS

### Animals and transgenic line generation

Mouse lines were generated by zygotic injection of Cas9-gRNA ribonucleopreoteins. Candidate sgRNAs were designed using sgRNA Scorer 2.0 (Chari et al., 2017) and subsequently tested for editing activity in P19 cells using an approach previously described (Gooden et al., 2021). The most potent candidates were then procured as synthetically modified RNAs (Synthego). Cas9 protein was generated in house (Protein Expression Lab, Frederick National Lab) using an E. coli expression plasmid obtained as a gift from Niels Geijsen (Addgene #62731) (D’Astolfo et al., 2015). Superovulated C57Bl6NCr female mice were used as embryo donors. Zygotes were collected on day E0.5, microinjected and allowed to recover for 2 hours by incubation (5% CO2, 37°C), following which, viable embryos were surgically transferred to oviducts of pseudopregnant recipient females. The microinjection cocktail comprised of 75 ng/µl of in vitro synthetized gRNAs (Synthego) and 50ng/µl of Cas9 protein (produced by the Protein Expression Laboratory, Frederick National Lab) in 50µl total volume, kept on dry ice until just prior to microinjection. For insertions requiring repair templates, 75 ng/µl of single-stranded DNA was added to microinjection cocktail. Founder mice were then bred to C57Bl6 mice obtained from The Jackson Laboratory. The CTCFΔ(2-4) and SCRΔ lines were generated using 2 gRNAs to achieve a deletion. The CTCFΔ(C5) line was obtained by replacing the CTCF 19bp motif by a sequence of the same size containing a NotI restriction enzyme site for easier genotyping. For this line, 2 gRNAs were used to generate two DNA double stranded breaks. The CTCFi3X(-) and CTCFi3X(+) lines were generated by inserting three consecutive CTCF motifs (Redolfi et al., 2019) in two different locations between Sox2 and SCR. For this, one gRNA for each location was used to induce double-stranded DNA breaks and repair templates containing the CTCF motifs flanked by homology arms were used for zygotic injection. The CTCFi18x(+) line was a byproduct of injection of the CTCFi3X(+) repair template where one chimeric F0 mouse was obtained containing three alleles: wild-type, CTCFi3X(+) and CTCFi18x(+). This founder male was bred to C57Bl6 females and heterozygous F1 progeny for each of the three alleles were recovered at similar levels. To identify the sequence of the integration of the CTCFi18x(+) line we amplified gDNA from ES cells derived from homozygous blastocysts using a primer pair surrounding the insertion sites. This large, repetitive DNA fragment was then sequenced using Pacbio, which revealed 6 concatamers of the repair template for a total of 18 CTCF motifs all in the same orientation. The two lines, CTCFi3X(+) and CTCFi18x(+), were then crossed separately. For generation of CTCFi3x(-):3X(+), homozygous females of the CTCFi3x(-) were used as female donors and zygotes were injected with the CTCFi3X(+) repair template and gRNA-Cas9 ribonucleoprotein. Repair templates, gRNAs, primers used for genotyping, and sequencing of ligation junctions of the deletion lines can be found in Table S1. In addition, the Sox2-GFP line was purchased from the Jackson Labs (strain 017592) (Arnold et al., 2011). All mouse studies were performed according to NIH and PHS guidelines and only after protocols were approved by the Animal Care and Use Committees of the National Cancer Institute and Eunice Kennedy Shriver National Institute of Child Health and Human Development.

### ES cell line establishment

Blastocysts derived from natural breeding of heterozygous transgenic mice were used for ES cell line establishment following a published protocol with some modifications (Czechanski et al., 2014). Presence of copulatory plug in the early morning was designated as E0.5. Blastocysts at E3.5 were harvested by flushing female uterine horns with M2 medium. 24hrs prior to embryo isolation, primary mouse embryonic fibroblasts (MEFs) were thawed and grown in 24 well plates in MEF culture medium containing Dulbecco’s Modified Eagle Medium (DMEM) (Thermo Fisher, #11965118), 10% FBS (VWR, #97068-091), 2 mM Glutamax (Thermo Fisher, #35050079) and penicillin–streptomycin (Thermo Fisher, #15140163). Isolated blastocysts were plated over MEFs in presence of serum free media combined with MEK/ERK pathway inhibitor (PD0325901, Reprocell, #04-0006-02), GSK3 signaling inhibitor (CHIR99021, Reprocell, #04-0004-02) and leukemia inhibition factor (LIF, Sigma, #ESG1107). Serum free media comprised of Neurobasal medium (Thermo Fisher, 21103049), DMEM/F12 Nutrient mixture (Thermo Fisher, 21103049), 1% penicillin– streptomycin (Thermo Fisher, #15140163), 2 mM Glutamax (Thermo Fisher, #35050079) and β-mercaptoethanol supplemented with N2 (Thermo Fisher, #17502001) and B-27 (Thermo Fisher, #17504001). Single blastocysts, plated in individual wells were cultured over feeders for 10 days at 37oC, 5% CO2 to allow ICM outgrowth and 50% of the medium was changed every 48 hrs. Embryos hatched out and attached to the feeder layer within 48-72 hrs of plating. Outgrowths were picked up using a mouth pipette without feeder carry-over and transferred into 96 well round bottom plates containing 50 µl of cold 0.05% trypsin and incubated for 5 minutes at 37oC. The outgrowth was disaggregated into small clumps of cells by pipetting up and down vigorously several times while observing under the microscope. Trypsin was quickly inactivated by adding 250 µl of 20% serum containing medium and LIF. From this cell suspension, 100 µl were lysed and used for genotyping and the remaining 200 µl were plated in 96 well plates containing feeders and grown for 48-72hrs in presence of ES serum-containing media until ES colonies were visualized. Meanwhile genotype of each ES clone was determined, and wildtype and homozygous clones were propagated for additional 2-3 passages in presence of feeders and serum containing media. At least two independent cell lines from two individual blastocysts were established for each genotype. The growth and morphology of each ES clone was monitored before freezing the stocks (P3 or P4). Serum containing media comprised of Knockout Dulbecco’s Modified Eagle Medium (DMEM) (Thermo Fisher, #10829018) with 15% FBS (VWR, #97068-091), 2 mM Glutamax (Thermo Fisher, #35050079), 0.1 mM β-mercaptoethanol, 0.1 mM MEM non-essential amino acids (Thermo Fisher, #11140050), 1 mM sodium pyruvate (Thermo Fisher, #11360070), 1% penicillin–streptomycin (Thermo Fisher, #15140163) and Recombinant mouse LIF (Sigma, #ESG1107). Additionally, a fraction of these ES cell clones were feeder depleted and cultured in serum free media (2i + LIF) for at least 4-5 passages and frozen in freezing medium (95% FBS + 5% DMSO) once they adopted a round 3D morphology and good growth rate. For all our downstream experiments early passages of frozen stocks of ES cell clones were used without feeders grown in serum free media in the presence of 2i + LIF to mimic as close as possible the in vivo pluripotent epiblast of blastocysts.

### RNA quantification of *Sox2* expression

*Sox2* expression was quantified in embryos (E3.5 blastocysts and E11.5 heads) and ES cells. Isolation of DNA and RNA from single blastocysts was performed as previously described (Huffman et al., 2012) with a few modifications. Reagents from Dynabeads mRNA DIRECT Purification Kit (Thermo Fisher, 61012) were used. Briefly, late-stage blastocysts at E3.5 were flushed out of uterine horns of super-ovulated females using M2 media and transferred into 50 µl of pre-warmed lysis buffer (100mM Tris-HCl pH-7.5, 500mM LiCl, 10mM EDTA pH-8, 1%LiDS, 5mM DTT) in 0.2 ml DNA low binding PCR tubes. Lysed embryos were stored in -20 and processed within a week. Dynabeads Oligo(dT)25 mRNA isolation beads were warmed at room temperature for 30 mins and rinsed with 100 µl of lysis buffer by vortexing continuously for 5 mins. 10 µl of bead suspension was used per 50 µl of thawed out embryo lysate. The poly-A tail of mRNA was allowed to anneal to beads by mixing the embryo lysates and bead suspension in a vortexer for 5 mins at low speed, followed by 5 mins incubation without shaking. Tubes were briefly spun and placed in a magnetic stand to collect the clear supernatant, which was used for DNA isolation using 2X SPRI beads to determine genotype. mRNA bead complexes were washed twice using Wash Buffer A and twice with Wash Buffer B by vortexing for 5 mins each. In the final wash step, buffer was replaced with 10 µl of first strand synthesis mix (1X SSIV Buffer with 0.5mM dNTP, 5mM DTT, 20U of RNase Inhibitor and 20U of SuperScript reverse transcriptase, Thermo Fisher, #18090010) to resuspend the beads and prepare cDNA directly. The mixture was incubated at 50oC for 20 mins and flicked every 5 mins to make sure that beads are in suspension. Enzyme inactivation was done on a PCR machine at 95oC for 1 min and cDNA was collected quickly from the beads and frozen until use. *Sox2* and *Gapdh* levels were quantified by qPCR using iTaq Universal SYBR Green Super mix (Biorad) in a Roche Light Cycler (Conditions include an initial holding stage at 95oC for 30s followed by 40 cycles at 95oC for 15s and 60oC for 1 min). E11.5 head tissue was dissected in cold PBS and lysed in 1 ml of TRIzol reagent (Thermo Fisher). Tissues were homogenized at room temperature by passing it a few times through a syringe with a 23G needle (BD Biosciences). ES cell RNA was also isolated by directly lysing cell pellets in TRIzol reagent. Following a chloroform step, RNA in aqueous phase was precipitated with isopropanol, followed by ethanol wash, and resuspended in water. RNA was quantified using Qubit and subsequently used for cDNA synthesis according to user manual for SuperScript reverse transcriptase (Thermo #18090010). Fold change in *Sox2* expression in mutants compared to wild type littermates or ES clones was calculated using the ΔΔCT method and each sample was normalized to *Gapdh* levels. Primers used for qPCR are listed Table S1.

### Immunofluorescence

Mouse embryos at different developmental stages used for immunostaining were collected in cold PBS solution using standard dissection protocols (Behringer R, 2014). Blastocysts were fixed with 4% formaldehyde at room temperature for 10 mins. Unhatched embryos with intact zona pellucida were digested with Acid Tyrode solution prior to fixation. For E6.5 and E7.5, incubation time was 30 mins and for older embryos and tissues, overnight fixation at 4°C was used. Next, embryos were washed thrice with PBX (0.1% Triton X-100 in PBS) and permeabilized with 0.5% Triton X-100 and 100mM Glycine in PBS for 30 mins at room temperature. E9.5 embryos and tissues required additional steps such as gradual dehydration into 100% methanol, overnight bleaching (one part 30% hydrogen peroxide and 2 parts of methanol), and Dents fixative (20% DMSO in methanol at 4°C). Preceding antigen blocking, E9.5 embryos and tissues were re-transferred into PBST (0.1% Tween-20 in PBS) gradually by a series of decreasing methanol concentrations. Embryos were incubated with blocking solution (2% horse serum in PBS for blastocysts, 5% horse serum in 0.2% BSA for E6.5/7.5 and 10% FBS in PBST for E9.5 and tissues), followed by primary antibody incubation overnight at 4°C. For E9.5 embryos and tissues incubation time was for 72 hrs. Embryos were then washed with PBX, incubated in blocking solution, and transferred into secondary antibodies in blocking buffer. Blastocysts were incubated at 4°C for 2hrs, E6.5/7.5 at room temperature for 2hrs and E9.5 and tissues at 4°C for 48hrs. Following washes in PBX specimens were counterstained with Hoechst solution (1:5000 in 0.1% PBX) and washed again and imaged. Confocal images of blastocysts, E6.5 and E9.5 were captured using a Zeiss LSM880 laser-scanning microscope. Image analysis and quantification of protein expression was performed in FIJI (Schindelin et al., 2012). Normalization of signal intensity between litters was done by dividing the intensity value of each cell by the highest value recorded in heterozygous littermates. Images of E9.5 and AFG tissues were captured in a Leica M165C dissection microscope. In all imaging steps, embryos of the same litters were processed in parallel and imaged with the same conditions.

### Capture Hi-C

Hi-C libraries for capture Hi-C were prepared as described in (Thompson et al., 2021). Briefly, for mouse ES cell lines, 1 million cells per sample were trypsinized, washed in growth media and fixed for 10 minutes at room temperature while rotating with 1% formaldehyde (Thermo: 28908) in 1ml of HBSS media. CHiC was processed separately for the two independent lines of each genotype that had been established from two independent blastocysts. Heads of individual E11.5 embryos consisting of forebrain, midbrain and hindbrain were dissected in PBS with 10% FBS. After cutting specimens into small pieces with fine scissors, samples were incubated for 45 mins at 37°C in 300µl of 90%FBS/0.1mg/ml Collagenase I (Sigma:CO130) in PBS. Following incubation, material was strained through 70µm filter and washed with cold PBS before incubating for 10 mins at RT in 1% formaldehyde. At least two different embryos pergenotype were used as separate replicates for these experiments. For isolation of AFG-derived tissues from Sox2-GFP heterozygous embryos, E11-E12.5 embryos were dissected in PBS with 10% FBS to microscopically dissect the lung and stomach primordia as well as the trachea and esophagus. GFP+ cells from non-AFG tissues, such as neural tube cells were not carefully removed during dissection. These samples were then cut into small pieces and incubated for 45 minutes at 37 °C while shaking in a buffer containing: Collagenase IV (2.2 mg/mL; Gibco), BSA (0.1%; Jackson Laboratory), DNase (125 units/ mL; Worthington), HEPES (20 mM), CaCl2 (1 mM), Pluronics (1%; (p-188 aka F-68; Thermo-Fisher) in Medium 199 (Gibco) (Zalc et al., 2021). Samples were then sorted for GFP+ population using Hoechst staining to isolate living cells. As control for setting gates, material isolated and processed in parallel from wt littermates was used. GFP+ cells were pooled from different litters containing cells isolated from several Sox2-GFP heterozygous embryos. Following FACS, samples were fixed as described above. To stop fixation, Glycine was added at final concentration of 0.13M and incubated for 5 minutes at RT and 15 minutes on ice. Cells were then washed once in cold PBS, centrifuged at 2500g 4°C for 5 mins (these centrifugation conditions were used for all washes following fixation) and pellets frozen at -80°C. Thawed cell pellets were incubated in 1ml lysis buffer (10mM Tris-HCL pH8, 10mM NaCl, 0.2% Igepal CA-630, Roche Complete EDTA-free Sigma #11836170001). Following lysis, cells were dounced for a total of 40 strokes with a “tight pestle” and then washed in cold PBS. For DpnII digest, cells were resuspended in 50µl 0.5% SDS and incubated at 62°C for 10 minutes. Then 150µl of 1.5% Triton-X was added and cells incubated for 15 minutes at 37°C while shaking at 900rpm. 25µl of 10X DpnII restriction buffer (NEB) was added, and cells further incubated for 15 minutes while shaking. 200U of DpnII (NEB R0543M) were then added and incubated for 2h, then 200U more and incubated overnight. Next morning 200U more were added and incubated for 3h (total 600U of DpnII). DpnII was inactivated at 62°C for 20 minutes. Biotin fill-in was done by incubating cells with a mixture of 4.5 µl dCTP dTTP and dGTP at 3.3 mM, 8µl klenow polymerase (NEB M0210L) and 37.5µl Biotin-14-dATP (Thermo 19524016) for 4h at RT while shaking at 900rpm for 10 seconds every 5 minutes. Ligation was done overnight at 16°C also rotating at 900rpm for 10 seconds every 5 minutes by adding 120µl of 10X ligation buffer (NEB), 664µl water, 100µl 10% Triton-X, 6µl BSA 20mg/ ml, and 2µl T4 ligase (NEB cat #M0202M). Crosslink removal was done overnight with 50µl of proteinase K in 300µl of following buffer (10mM Tris-HCl pH8.0, 0.5M NaCl, 1%SDS) while shaking at 1400rpm at 65°C. Following Sodium Acetate and 100% Ethanol -80°C precipitation, DNA was resuspended in 50µl 10mM Tris HCL. Sonication was done using Covaris onetube-10 AFA strips using the following parameters for a 300bp fragment size (Duration: 10secs, repeat for 12 times, total time 120 secs, peak power-20W, duty factor 40%, CPB-50) in a Covaris ME220 sonicator. Sonicated material was then size-selected using SPRI beads with the following ratios: 0.55X and 0.75X. Hi-C material was then bound to 150µl Streptavidin C1 beads (Thermo 65002), washed and recovered following manufacturers recommendations. Bead-bound DNA was resuspended in 50µl 10mM Tris HCl. Library preparation was done using the Kapa Hyper Prep KK8502 kit. 10µl of End-repair buffer and enzyme mix were added to resuspended beads and incubated for 30 minutes at RT and then 30 minutes at 65°C. 1µl of 15mM annealed-Illumina adaptors, containing a universal p5 and an indexed p7 oligo, were then incubated with a mixture containing 40µl of ligase and ligation buffer at RT for 60 minutes. Libraries were then amplified using 4 reactions per sample for a total of 200µl and 10 cycles, as recommended by manufacturer. For capture, 1µg of Hi-C library per sample was mixed with 5 μl of SureSelect XT HS and XT Low Input Blocker Mix (Agilent). Samples were denatured at 95°C for 5 minutes and pre-hybridized for 10 minutes at 65°C. 2 μl of SureSelect probes (Agilent) were mixed with 2 μl of 25% RNAse block, 6 μl of Hybridization buffer and 3 μl water. Following pre-hybridization, this probe mixture was added to pre-hybridized samples and incubated for 1 hour at 65 °C. For washing, samples and probes were bound to 50 μl Streptavidin C1 beads and washed using SureSelect wash buffers as recommended by the manufacturer. Washed material was resuspended in 25 μl of water and amplified by PCR using using 20 cycles.

### Capture Probe Design

RNA probes were designed using GOPHER (github.com/TheJacksonLaboratory/Gopher) and the following options: input region (mm10) chr3: 34016000-35655100, extended approach, maximum kmer alignability of 1, margin size 280, GC 35/65. For output we used minimum probe count and unbalanced margins. GOPHER output coordinates were then used as input for SureSelect probe design tool (Agilent) with maximum masking stringency and balanced boosting and 1x tiling density. Regions covered by biotinylated RNA probes can be found in **Table S2**.

### Capture Hi-C analysis

CHiC libraries were sequenced with paired-end reads of 51 nucleotides. Data was processed using the Hi-Cpro pipeline (Servant et al., 2015) to produce a list of valid interactions pairs with the Capture Hi-C option for the following region: chr3:34016000-35655100. This list was converted into cool and mcool files for visualization with higlass (Kerpedjiev et al., 2018). Following assessment of replicate reproducibility, the replicates of same condition were merged for visualization using Hi-Cpro followed again by conversion to cool and mcool files. 1D Capture Hi-C heatmaps are shown at 4kb resolution and 2D heatmaps for zoom ins at 2kb. Signal across all samples was normalized according to signal in WT samples. Information on number of reads and peaks can be found in Table S2. Differential 1D Capture Hi-C heatmaps were produced using the divisor function of higlass and signal from mutant lines was divided by WT Capture Hi-C. Heatmap values were then normalized to the sample with highest differential scores. To calculate insulation scores we used FANC (Kruse et al., 2020) using 5kb Capture Hi-C resolution and a 50 kb sliding window. The make_viewpoints Hi-Cpro script was used to obtain virtual 4C plots for each replicate of each viewpoint. The following regions were used as viewpoints (Sox2 -chr3:34647939-34653212, SCR - chr3:34752710-34760133, DNE - chr3:35083629-35091662). For visualization, averages of each replicate were used and signal was normalized with DE-Seq2 using 5kb overlapping windows by sliding each window 1kb. p-values were calculated using DESeq2 and comparing Capture-C signal of 2 replicates over the regions shown in figure. All samples were compared to WT. Adjusted p values lower than 0.01 were considered as threshold for significance and are shown above the plot. Zoom in for Capture HiC data of E11.5 is shown using combined replicate signal at fragment level.

### ChIP-seq

Heads of E11.5 WT embryos were dissected as described above for CHiC. Tissues were then processed following Encode guidelines as described in (Gorkin et al., 2020). Following dissection, samples were quickly pulverized with liquid nitrogen, dissociated with mortar and pestle and transferred to 10ml of cold PBS. 1ml of crosslinking buffer (0.1M NaCl, 1mM EDTA, 0.5mM EGTA, 50mM HEPES pH8.0, 11% formaldehyde) was added and samples were fixed for 20mins at RT followed by neutralization with 1 ml of 1.4M Glycine. Following centrifugation at low speed, supernatant was removed, and cells were used for immuno precipitation. Pellets were lysed in Farnham lysis buffer (5mM PIPES PH 8.0, 85mM KCl, 0.5% NP-40, supplemented with protease inhibitors). To lyse cells, pellets were reconstituted in 1ml buffer/ 10 million cells and incubated on ice for 10 mins, following which they were dounced and then centrifuged at 2000g, at 40C for 5 mins. The pelleted nuclei were then resuspended in 500µl RIPA (50 mM Tris-HCl pH 8.0, 150 mM NaCl, 2 mM EDTA pH8, 1% NP-40, 0.5% Sodium Deoxycholate, 0.1% SDS, supplemented with ROCHE Complete protease inhibitor tablets (no EDTA), mixed gently by pipetting, and incubated on ice for 10 mins to achieve complete lysis. Extracted chromatin was then sonicated using Covaris milliTUBE 1ml AFA using the following parameters (Total duration: 40 mins, peak power-20W, duty factor 5%, CPB-200, paused every 10 mins) in a Covaris ME220 sonicator. Sonicated material was then spun down at 20000g, at 40C for 15 mins. Supernatant was collected in fresh tubes and chromatin was quantified using Qubit high sensitivity DNA kit. 30µg of chromatin was used for the ChIP reactions in a total reaction volume of 1ml, adjusted with RIPA. 30 µl of Protein A/G beads pre-conjugated to 2 µg antibody (by incubation of antibody with washed beads resuspended in 500µl RIPA for 3h) was added to the reaction and chromatin was capture by overnight incubation at 40C, on a rotating platform. The next day, captured chromatin was washed successively as follows: once in low-salt wash buffer (0.1% SDS, 1% Triton X-100, 2 mM EDTA, 20 mM Tris-HCl pH 8.0, 150 mM NaCl), twice in high-salt wash buffer (0.1% SDS, 1% Triton X-100, 2 mM EDTA, 20 mM Tris-HCl pH 8.0, 500 mM NaCl), twice in lithium chloride wash buffer (250mM LiCl, 1% NP-40, 1% Sodium Deoxycholate, 1 mM EDTA, 10 mM Tris-HCl pH 8.0), and twice in 1X TE (10 mM Tris-HCl pH 8.0, 1 mM EDTA). Bound chromatin was then eluted in 200 µl freshly prepared direct elution buffer (10mM Tris-HCl pH8, 0.3M NaCl, 5mM EDTA pH8, 0.5% SDS) supplemented with 2 µl RNAse A (10 mg/ml), by incubating on a thermomixer at 650C, at 800 rpm, overnight. The next day, the beads were discarded, and the supernatant was incubated with 3 µl of Proteinase K (20mg/ml) at 550C, 1200rpm, for 2h to reverse crosslinks. DNA was then eluted in 20 µl deionized water using the Zymo DNA Clean and Concentrator kit (Zymo, D4033). 10 µl of eluted DNA was used for library preparation exactly as described for CUT&RUN. 12 amplification cycles were used per sample, and library was sequenced on HiSeq2500 using PE50. Reads were processed with Bowtie2 (Langmead and Salzberg, 2012), with the following parameters (-N 1 –local --very-sensitive-local --no-unal --no-mixed --no-discordant --phred33 -I 10 -X 700 -x). Reads that mapped to ENCODE mm10 blacklist regions were removed using Samtools (Li et al., 2009). Piccard (broadinstitute.github.io/picard/) was used to remove duplicated reads. For visualization we used Deeptools to generate coverage profiles. Stats for number of reads can be found in Table S2.

## DATA ACCESSIBILITY

A list of publicly available data used in this study can be found in Table S2 which include data from the following studies (Bonev et al., 2017; Gorkin et al., 2020; Hansen et al., 2017; Kagey et al., 2010; Li et al., 2018; Thompson et al., 2021; Whyte et al., 2013; Xiang et al., 2020). All datasets were shown using processed files as available except for files mapped to mm9, which were converted to mm10 using CrossMap (Zhao et al., 2014). FASTQ and processed CHiC and RAD21 E11.5 head ChIP-seq data can be found in GEO under accession number GSE190359. CHiC data can also be easily navigated at: resgen.io/pedrorocha/sox2/views/

## ACKNOWLEDGMENTS

We would like to thank all members of the Unit on Genome Structure and Regulation for comments and discussions on this project and manuscript as well as Karl Pfeifer, Todd Macfarlan, and Judith Kassis. We thank tips by Heinrich Schrewe for AFG-isolation and Nestor Saiz for blastocyst IF and image analysis. We thank NICHD’s molecular genetics core, specifically Steven Coon, Tianwei Li and James Iben. This work utilized the computational resources of the NIH HPC Biowulf cluster (hpc.nih.gov). We thank Lisa Price for cell sorting experiments. We thank the mouse core of NICHD specifically Jeanne Yimdjo, Victoria Gibbs and Alexander Grinberg. We also thank the NCI’s molecular histopathology core, especially Tamara Morgan, Jennifer Mata and Baktiar Karim. This work was funded by NIH intramural project HD008975-02.

## AUTHOR CONTRIBUTIONS

SC, NK, AE and PR performed experiments. PA and RC designed and generated transgenic mouse lines. SC, NK and PR analyzed data. SC and PR wrote manuscript with input from all the authors.

---

**Supplementary figure S1:**
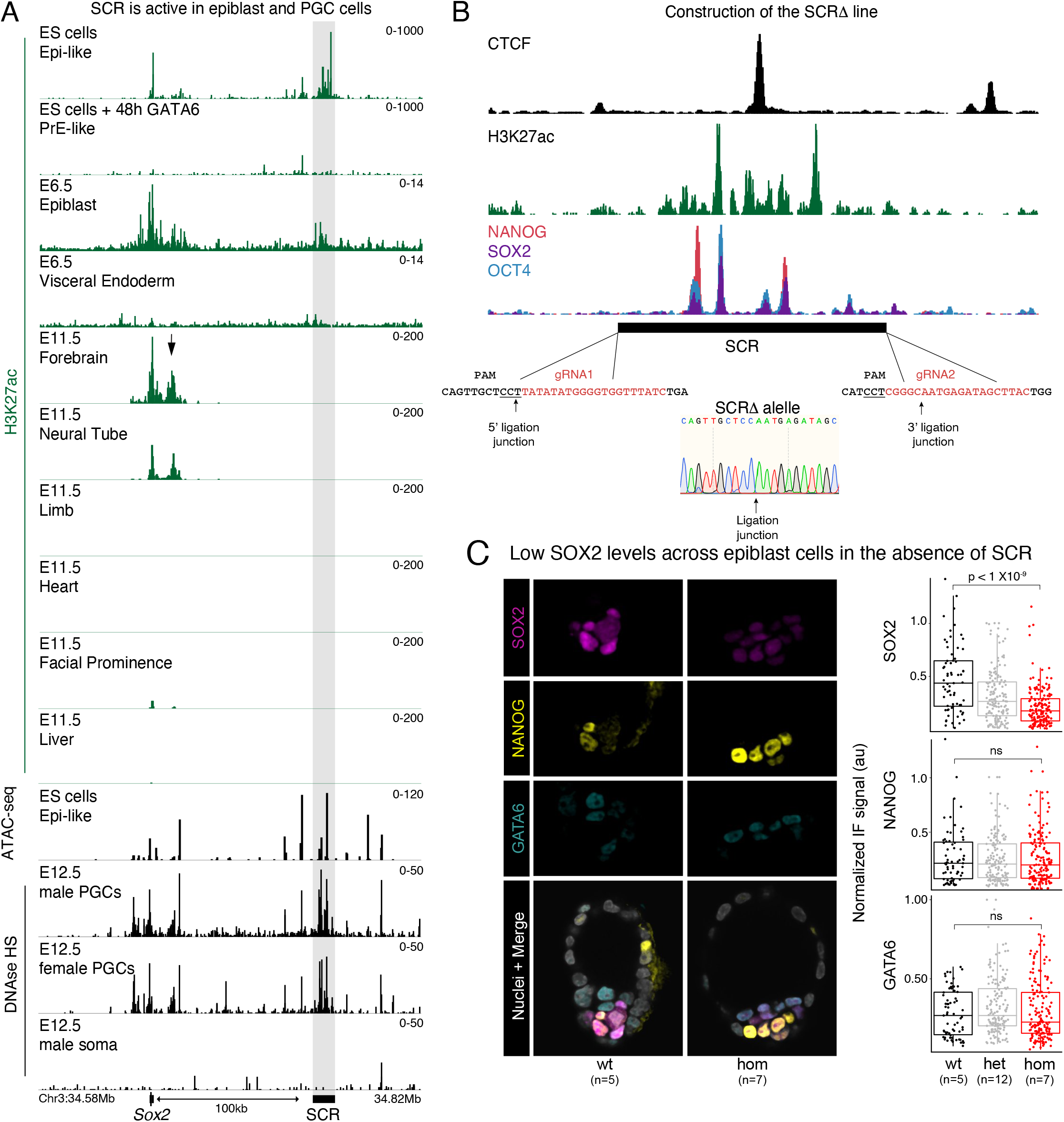
SCR is required for *Sox2* expression. **A** *Sox2* and SCR activity during early mouse development as assessed by enrichment of H3K27ac and ATAC-seq. H3K27ac CUT&RUN in mES cells cultured epiblast-like conditions shows enrichment at both *Sox2* and SCR but not in cells differentiated into primitive endoderm (PrE). No H3K27ac is detected at *Sox2* or SCR in the visceral endoderm of post-implantation embryos and SCR activity is diminished in epiblast cells. Despite strong enrichment at the *Sox2* gene body in the neural tube and forebrain of E11.5 embryos, SCR is no longer active. Instead, other proximal regions (black arrow) show increased activity. ATAC-seq data shows that both male and female primordial germ cells, but not the surrounding soma, are enriched in accessible chromatin at SCR. See Table S2 for sources on publicly available data used to assemble this panel. **B** Region targeted for deletion in the SCRΔΔline and gRNAs used. Browser shot shows enrichment of CTCF, NANOG, SOX2, OCT4 and H3K27ac over *Sox2* and SCR in ES cells. Nucleotides in red represent the protospacer sequence of gRNAs used for injection. Underlined nucleotides highlight location of the Cas9 PAM in the mouse genome. Arrows on both sides of the deletion highlight the ligation junction detected in the mouse used as founder as determined by Sanger sequencing. **C** IF of E4.5 blastocysts stained with antibodies targeting GATA6, NANOG and SOX2. Each dot represents a cell and to allow comparison across three different litters the intensity of each cell was normalized by the cell with highest intensity in heterozygous embryos. Number of embryos analyzed for each genotype is shown below the plot. A Wilcoxon two-sided test was performed to assess statistical significance.

**Supplementary figure S2:**
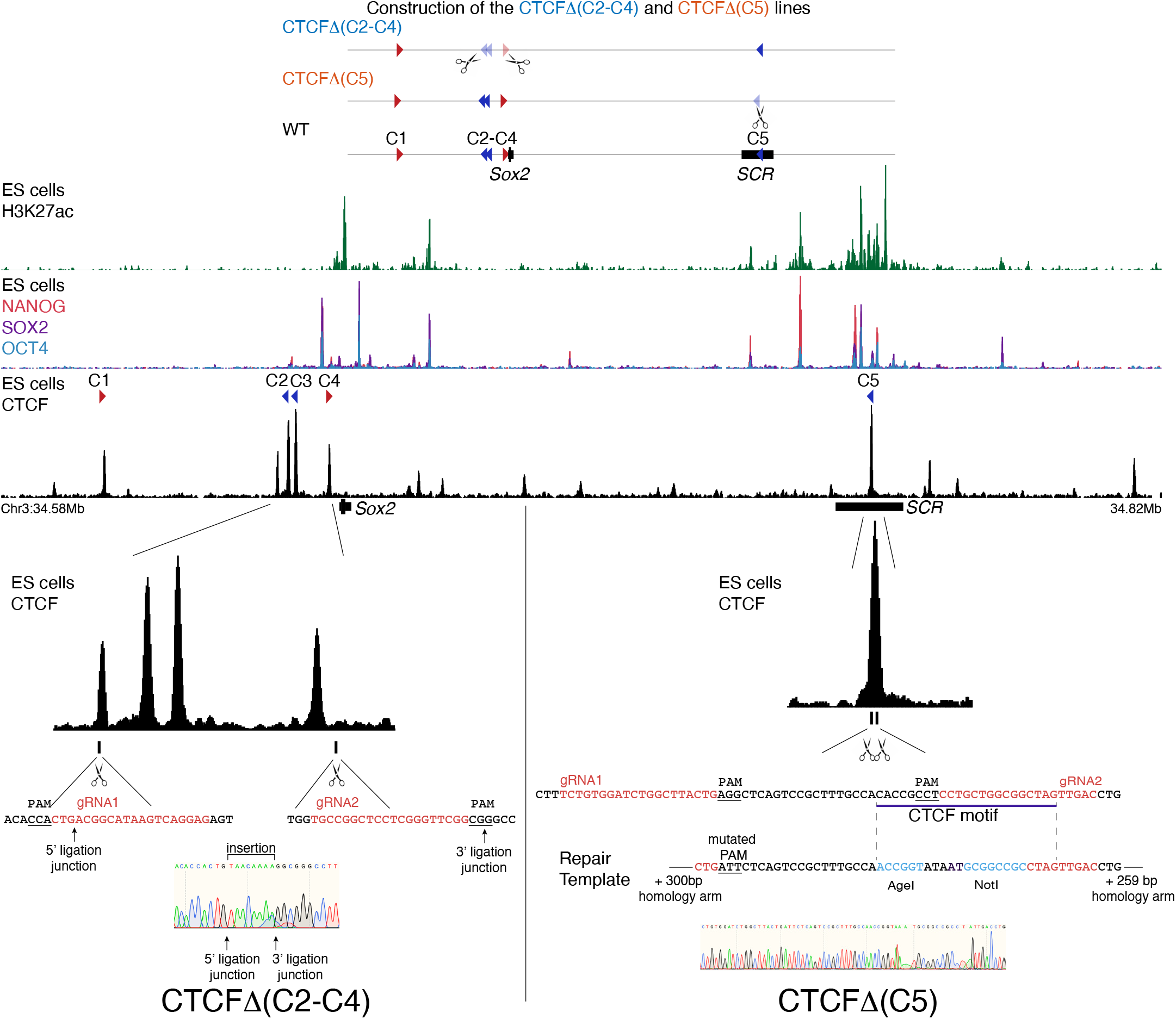
Construction of the CTCFΔ(C2-C4) and CTCFΔ(C5) lines. Scheme depicting targeting strategy for generation of the two transgenic lines. Browser shot shows enrichment of CTCF, NANOG, SOX2, OCT4 and H3K27ac over Sox2 and SCR. A NANOG/SOX2/OCT4 peak showing high enrichment was deleted in the CTCFΔ(C2-C4) but this region has been previously shown to not be required for mouse development. Browser shots with zoomed-in views of CTCF enrichment at *Sox2* and SCR show precise location of gRNAs used in zygotic injections. CTCF peak nearest the most centromeric gRNA used in the CTCFΔ(C2-C4) targeting does not contain a significant CTCF motif according to FIMO. However, this peak was still deleted in the CTCFΔ(C2-C4) line. Nucleotides in red represent the protospacer sequence of gRNAs used for injection. Underlined nucleotides highlight location of the Cas9 PAM in the mouse genome. Arrows on both sides of the deletion highlight ligation junction detected in the mouse used as founder as determined by Sanger sequencing. For the CTCFΔ(C5) scheme, purple line highlights location of the CTCF motif at SCR. The central sequence of the repair template is shown, with parts of the protospacer sequence used in the gRNA shown in red, and the restriction enzyme target that replaced the CTCF motif in blue. Full sequence of homology arms on both sides is omitted.

**Supplementary figure S3:**
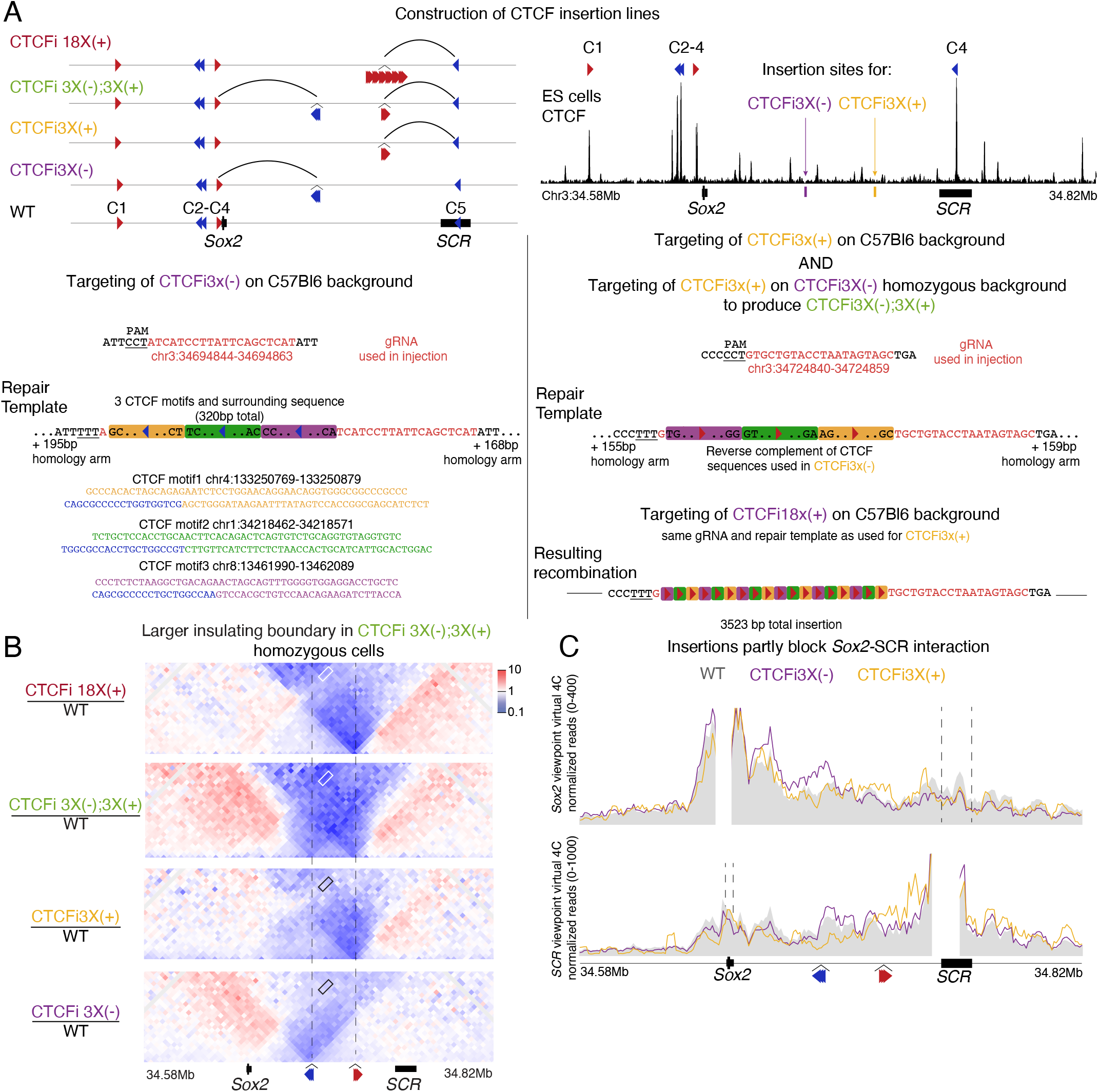
SCR can bypass strong insulation generated by CTCF-mediated loops. **A** Top left scheme depicts targeting strategy for generation of the transgenic lines carrying insertion of CTCF motifs. Arcs represent predicted CTCF-mediated loops generated in each insertion line as predicted by the loop-extrusion model. Top right browser shows CTCF ChIP-seq enrichment and insertion sites of CTCF transgenes. Left bottom panel shows the targeting of the C57Bl6 genome to generate CTCFi3x(-) mice. Right bottom panel shows targeting to generate CTCFi3x(+), CTCFi3x(+),CTCFi3x(-), and CTCFi18x(+) mice. Nucleotides shown in red represent the protospacer sequence of gRNAs used for injection while underlined nucleotides highlight location of the Cas9 PAM in the mouse mm10 genome. gRNA mm10 coordinates are shown in red. The central sequence of the repair templates is shown, with parts of the protospacer sequence used in the gRNA shown in red and mutated PAM nucleotides underlined. The complete sequence of homology arms on both sides is omitted. Each colored rectangle represents a different region from the mouse genome containing a CTCF motif and adjacent regions. Complete sequences of the CTCF motifs and adjacent regions are shown in different color with the central CTCF motif in blue representing targeting to negative strand in the CTCFi3x(-) line. The first and last two nucleotides of each of the three CTCF regions are shown in the schema of the repair template. The same three CTCF-carrying regions were used in CTCFi3x(-) and CTCFi3x(+) lines but targeted to different strands and locations. Therefore, the central region of repair template is the same but in different strands and containing different repair templates. Retargeting of the CTCFi3x(+) transgene on a homozygous CTCFi3x(-) background generated the CTCFi3x(-),CTCFi3x(+) line. The CTCFi18x(+) line was obtained as a consequence of the CTCFi3x(+) injection because of concatemerization of the repair template. The resulting allele is shown in the bottom right. **B** Differential CHiC interaction frequency heatmap. Red signal represents interactions occurring at higher frequency in mutant cell lines compared to control. Dotted lines represent insertion sites of CTCF transgenes. **C** Virtual 4C plot using *Sox2* and SCR viewpoints. Region surrounding viewpoint was removed from analysis. Dotted lines highlight SCR in the *Sox2* viewpoint (top), and *Sox2* in the SCR viewpoint (bottom). 4C signal is shown as average of the 2 replicates of each genotype in 5kb overlapping windows.

**Supplementary figure 4:**
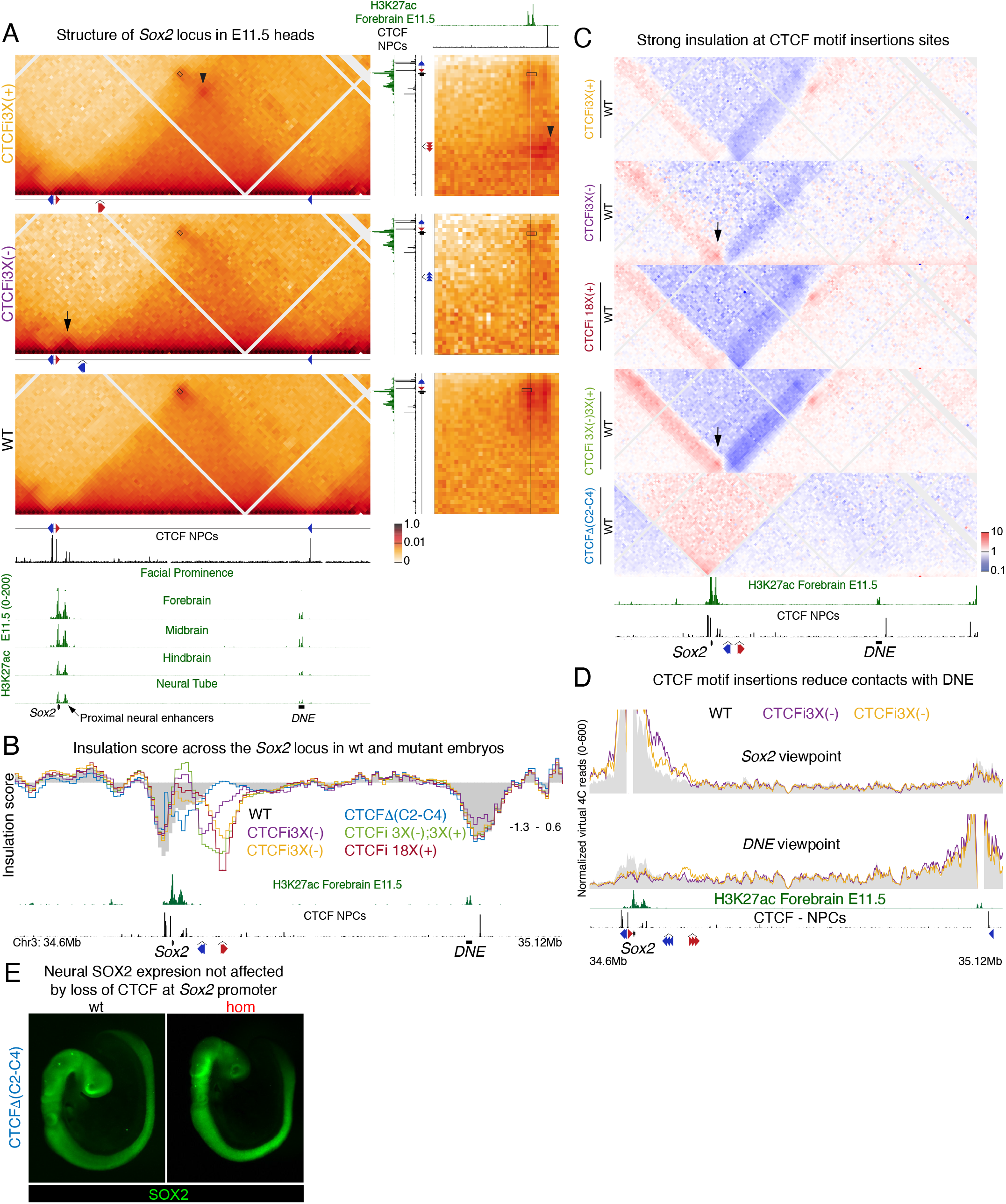
Interaction of Sox2 with long-distance neural enhancers bypasses CTCF-mediated insulation. **A** Interaction frequency heatmap determined by CHiC in heads of E11.5 embryos. Insets on the right show 2D interaction heatmaps highlighting interactions between regions surrounding *Sox2* and DNE. CTCF data shown under CHiC heatmaps from heads is derived from in vitro differentiated neural progenitor cells. Rectangles represent the *Sox2*-DNE interaction, arrowheads represent loops with CTCF downstream of DNE established by transgene insertions, black arrow represent loops with CTCF upstream of *Sox2* established by transgene insertions. **B** Insulation scores for 5kb windows in this region are shown where lower levels represent higher insulation. **C** Differential CHiC interaction frequency heatmap. Red signal represents interactions occurring at higher frequency in mutant cell lines compared to control. Black arrowhead highlights formation of a highly interacting domain containing *Sox2* and the proximal neural enhancers. **D** Virtual 4C plot using *Sox2* and DNE viewpoints. Region surrounding viewpoint was removed from analysis. E IF of E11.5 embryos stained with an antibody targeting SOX2.

**Supplementary figure 5:**
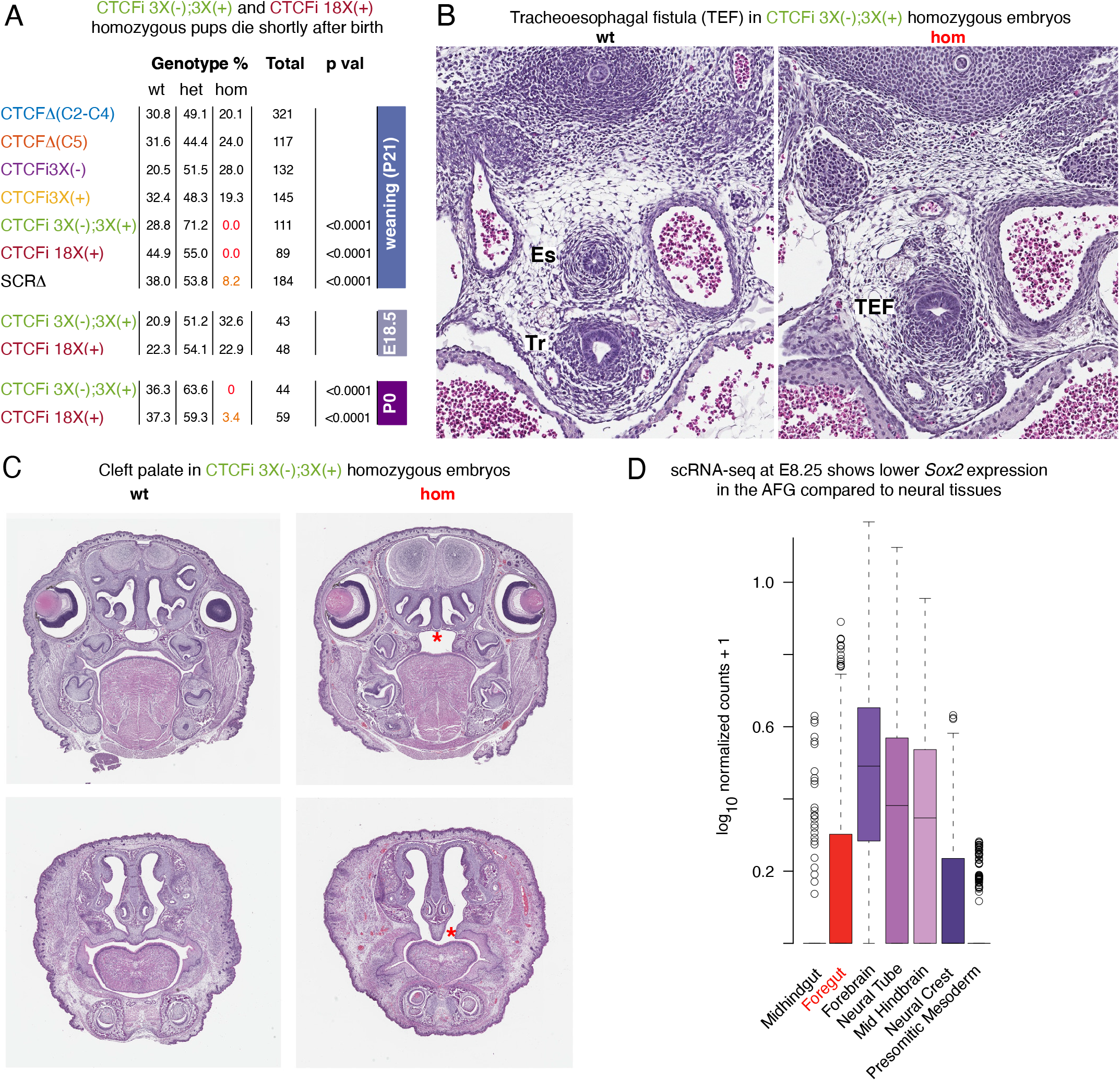
Developmental defects seen in homozygous embryos with modifications of the *Sox2* locus. **A** Genotyping at of living animals at weaning, P0 and E18.5 for the indicated strains. P values were calculated using a chi-squared test. B Transverse section of E13.5 CTCFi3x(-);3x(+) wt and homozygote littermates at heart level. Es-esophagus, Tr-trachea. **C** Frontal sections of E18.5 CTCFi3x(-);3x(+) wt and homozygote littermates. Asterisk highlights cleft palate defect. **D** Plot of normalized *Sox2* expression in single cells of WT E8.25 embryos. Plot was modified from marionilab.cruk.cam.ac.uk/organogenesis/

